# Rapid evolution and functional divergence of the monkeyflower *Mimulus lewisii* telomerase

**DOI:** 10.64898/2026.08.05.739867

**Authors:** Naseem Samo, Linh Nguyen, Surbhi Kumawat, Jae Young Choi

## Abstract

Telomeres are nucleoprotein structures that protect chromosome ends and are maintained by the Telomerase Reverse Transcriptase (TERT) protein that uses a noncoding Telomerase RNA (TR) as a template. In monkeyflowers, *Mimulus lewisii* had an ancient TR gene duplication, synthesizing an evolutionarily atypical sequence heterogeneous telomere. How TERT interacts with both TR paralogs during telomere maintenance is unknown and answers can shed novel insights underlying telomere function. Using new genome assemblies we discovered TERT is rapidly evolving in lineages sharing the TR duplication. We investigated the functional consequences arising from the rapid evolution, first by using yeast three-hybrid and testing the physical binding between conspecific and heterospecific TERT-TR combinations. Results showed TERT binds both ancestral (TR1) and derived (TR2) TR paralogs in *M. lewisii*, but not in species without a functioning TR2. We located the region of TR binding to amino acids near the KRxR motif. We then combined next-generation sequencing with Telomeric Repeat Amplification Protocol and discovered *M. lewisii* had high telomerase activity. Comparative transcriptomics indicated no strong evidence of expression divergence in telomere maintenance genes for *M. lewisii*, suggesting rapid evolution shaped TERT protein sequence. *In vivo* activity of *M. lewisii* telomerase was investigated by analyzing F1 telomeres generated by crossing *M. lewisii* and *M. verbenaceus*, which doesn’t have a functioning TR2. Results showed *M. verbenaceus* chromosome ends in the F1 had converted into *M. lewisii* telomeres, suggesting dominance of the *M. lewisii* telomerase. We demonstrate TERT-TR coevolution can have significant consequences on the evolution of plant telomeres.

**Significance statement:** Telomeres protect chromosome ends and are maintained by the telomerase complex. We discovered the catalytic component of the telomerase (TERT) was rapidly evolving in monkeyflowers (*Mimulus*) and studied the molecular consequences. In *M. lewisii,* TERT evolved lineage-specific amino acids to bind two sequence divergent telomerase RNA paralogs. Telomerase activity assay showed *M. lewisii* synthesized more telomere repeats compared to its sister species without the TR duplication, and transcriptomics indicated this was not due to a change in telomere maintenance gene expression. Genetic experiments in interspecies hybrids showed *M. lewisii* telomerase could convert chromosome ends in sister species into M. *lewisii*-like telomeres suggesting functional dominance. We show rapid evolution of the telomerase can have significant effects on telomere evolution.

## Introduction

A major goal of evolutionary research is to understand the functional consequence and evolutionary history of the amino acid differences between orthologous genes of different species. Neutral theory predicts the majority of the fixed differences have no fitness consequences (1), but the rare variants that do improve the organisms fitness will be selected by adaptive evolution (2). These adaptive changes have often resulted in functional and phenotypic novelty and many of these examples have become textbook cases of Darwinian evolution (3–5).

Proteins that need to interact with a multitude of molecules or respond to the environment (*e.g.* signal transduction, immunity, or enzyme catalysis) have been a hotspot for rapid evolution and adaptation (6, 7). This might suggest molecular systems with conserved core processes could be constrained from evolution and least likely to be targeted by adaptive evolution (8). Contrary to this expectation, however, in this study we provide evidence of rapid evolution occurring within the telomeres of monkeyflowers (*Mimulus*). The telomere is a G-rich DNA sequence that is found on the ends of almost all eukaryotic linear chromosomes and is critical for maintaining genomic stability (9, 10). Telomeres are bound by specialized proteins that protect chromosome ends from being detected as damaged DNA (9) and guard against attrition during cellular replication (11). Reflecting this functional importance, the sequence of the telomere is highly conserved in vertebrates where all lineages have a repetition of a TTAGGG sequence (12). Plants, on the other hand, display an exceptional diversity of telomere sequences (13, 14) suggesting the proteins that bind and maintain the telomeres might be rapidly evolving as well. For example, the single strand telomere DNA binding protein Protection of Telomeres 1 (POT1) has undergone an ancient gene duplication in Brassicaceae (15) and has undergone positive selection (16). Plant telomeres may hold unique examples of protein evolution relating to chromosomal maintenance and stability (17).

Previous studies have suggested the telomere sequence variation in plants was driven by the evolutionary divergence of the Telomerase RNA (TR) gene (18–22). During the maintenance of the telomere, the telomeric DNA is synthesized by the ribonucleoprotein enzyme complex called the telomerase, where the activity is mainly driven by the catalytic enzyme Telomerase Reverse Transcriptase (TERT) (23) and the noncoding RNA TR (24). The TR holds a templating sequence domain that is used by TERT for *de novo* synthesis of telomeric repeats at the 3′ ends of the telomere (25). Any sequence change at the TR templating sequence will be synthesized on the telomere DNA, hence the sequence evolution of the TR has been proposed to drive the sequence evolution of the telomere in plants (21, 22). A similar model has been proposed to underlie the telomere sequence variation in yeast as well (26).

We recently discovered the TR gene had duplicated in several species of *Mimulus* (22). *M. lewisii* in particular, belongs to the Erythranthe group (27) and harbors two TR paralogs. The ancestral TR paralog (termed TR1) was found in three other Erythranthe group species (*M. cardinalis*, *M. parishii*, and *M. verbenaceus*), while the newly derived paralog (termed TR2) was found in a subset of species with remnant sequences suggesting pseudogenization. The templating domain of TR1 encoded the sequence TTTCGG while TR2 encoded the sequence TTTCGGG (*i.e.* an additional G-nucleotide), indicating the duplication could result in a sequence heterogeneous telomere that consists of a mix of telomere repeats. The functional output of the telomerase was investigated using the Telomeric Repeat Amplification Protocol (TRAP) (28) and when applied to *M. lewisii* the TRAP products consisted of both TTTCGG and TTTCGGG repeats. In addition, nanopore long read sequencing analysis showed *M. lewisii* telomeres consisted of both TTTCGG and TTTCGGG repeat sequences. This indicated *M. lewisii* had a unique telomere maintenance mechanism where the telomerase used two TR paralogs to synthesize sequence heterogeneous telomeres. Several plant species also have TR duplications and their telomere sequences are a mix of different repeat sequences as well (21), suggesting the telomeres are an evolutionary dynamic compartment of plant chromosomes.

A puzzling question that arises from the *M. lewisii* telomeres is this: How does TERT bind with the two TR paralogs and generate a sequence heterogeneous telomere? There are elevated levels of sequence divergence between TR1 and TR2 (∼12% nucleotide differences), suggesting *M. lewisii* TERT must have evolved unique molecular changes to maintain the interaction with the two TR molecules. Additionally, it is unclear if because of the duplicated TR this would lead to a difference in telomerase activity for *M. lewisii* compared to the telomerase from other Erythranthe group species. In this study, we have addressed these questions by investigating the molecular evolution of *M. lewisii* TERT and its functional interaction with the TR paralogs. Molecular evolution analysis showed TERT was rapidly evolving in the Erythranthe group, and the molecular basis of this evolution was examined using yeast three-hybrid assay to test the interaction between conspecific and heterospecific TERT and TR combinations. Functions of the *M. lewisii* telomerase was studied by applying next generation sequencing with TRAP and comparing the telomerase activity across multiple species in the Erythranthe group. We also investigated if the transcriptional landscape had diverged between species with and without the TR duplication using RNA-seq. Lastly, we crossed *M. lewisii* with a sister species *M. verbenaceus* which does not have a duplicated TR, and telomere sequence analysis of the F1 suggested a telomerase dominance model where *M. verbenaceus* telomeres had converted into *M. lewisii* like telomeres. Our findings demonstrate that TERT has evolved the capacity to utilize multiple TR paralogs and has gained functions that suggest dominance during telomere maintenance.

## Results

### Telomere sequence variation within the Erythranthe group species

The Erythranthe group *Mimulus* comprises of 8 species (*M. cardinalis*, *M. cinnabarina*, *M. flammea*, *M. eastwoodiae*, *M. lewisii*, *M. parishii*, *M. rupestris*, and *M. verbenaceus*) (27) and previously we investigated the telomere sequence in 4 species (*M. cardinalis*, *M. lewisii*, *M. parishii*, and *M. verbenaceus*) (22). To gain a deeper understanding of the telomere sequence evolution within the Erythranthe group, we nanopore sequenced three additional Erythranthe group species (*M. cinnabarina*, *M. flammea*, and *M. rupestris*) and an outgroup species *M. platycalyx* from the Simiolus group. We generated between 6.7–21.1 Gbp of sequencing data, with read length N50 between 13.9–24.8 kbp, and mean read quality between 18.7–19.6 (Supplemental Table 1). Using the raw nanopore sequencing data we used the Topsicle package (29) to identify long reads sequenced from the telomere region and conduct sequence analysis using the telomere long reads. Results showed most Erythranthe group species had telomere long reads that were largely consisting of the 6 bp TTTCGG repeat sequence, while only *M. lewisii* had telomere long reads with a mix of 6 bp TTTCGG and 7bp TTTCGGG repeat sequences (Fig 1A). We then investigated if other ecotypes from *M. lewisii* also had a sequence heterogeneous telomere and nanopore sequenced three additional *M. lewisii* ecotypes. The sequencing output was similar to its sister species (Supplemental Table 1) and results showed all *M. lewisii* ecotypes had a sequence heterogeneous telomeres consisting of both 6 bp TTTCGG and 7bp TTTCGGG repeat sequences (Fig 1).

**Figure 1.**
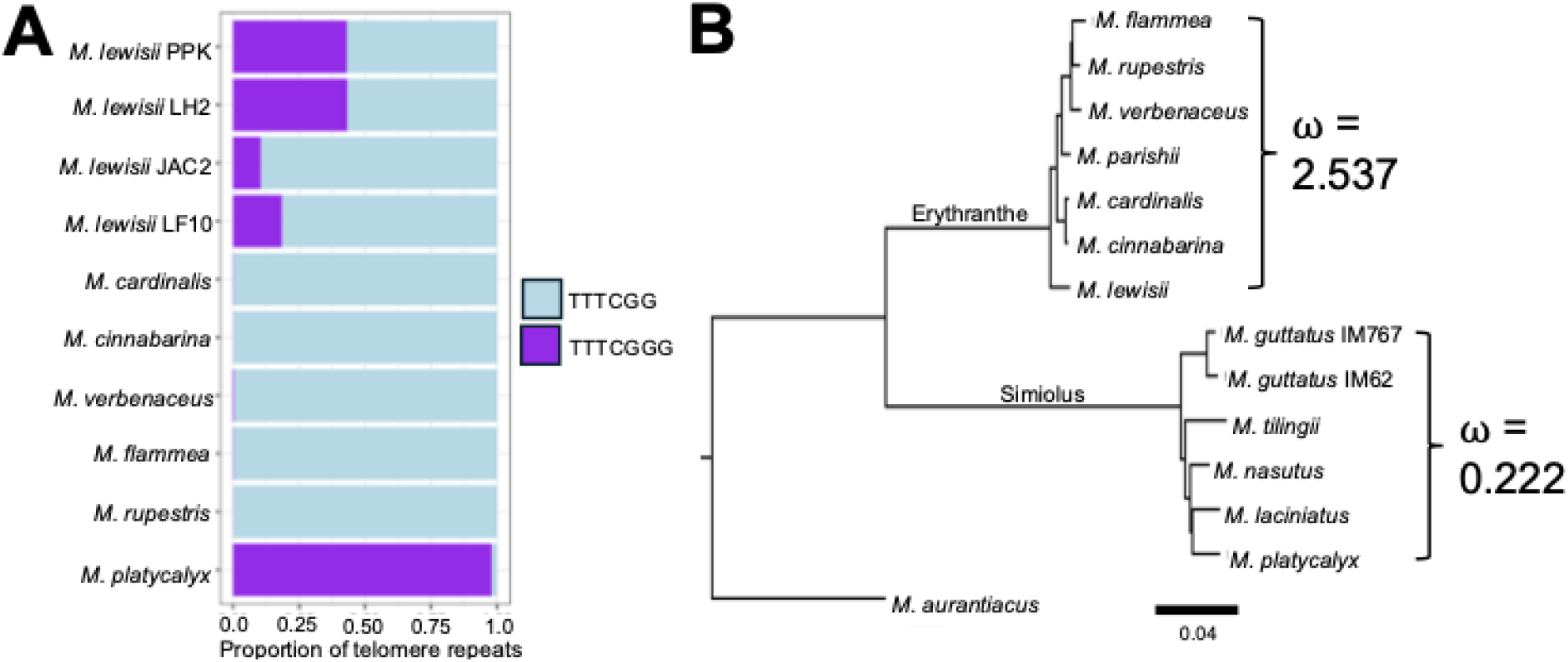
Evolutionary genomic analysis of *Mimulus* telomere and telomerase. (A) Proportion of long read sequences from the telomere that matches the 6bp TTTCGG repeat or 7 bp TTTCGGG repeat. (B) Phylogeny of TERT from multiple *Mimulus* species. The dN/dS ratio (ω) of the rapidly evolving site class was estimated using the PAML clade model (CmC) test.

### TERT is rapidly evolving in the Erythranthe group

The raw nanopore sequencing data from *M. cinnabarina*, *M. flammea*, and *M. rupestris* was used to generate *de novo* genome assemblies to conduct a comparative genomic analysis of TERT and TR in the Erythranthe group. The three species had genome assembly sizes over 440 Mbp and assembly N50 were between 660 kbp and 3.99 Mbp (Supplemental Table 2). Using the genome assemblies we searched for the TR2 gene using BLAST and discovered partial remnants in the *M. flammea* and *M. rupestris* genome assemblies (Supplemental Table 3). In *M. cinnabarina*, BLAST matched a full TR2 sequence but it is unlikely to be functional given its telomeres were enriched only for the 6 bp TTTCGG repeat (Fig 1A). This suggests the TR2 was potentially undergoing pseudogenization in *M. cinnabarina* similar to the TR2 gene in *M. cardinalis* (22). Based on previous findings and this study, seven out of the eight Erythranthe group species had a full or partial TR2 gene indicating the duplication was old and occurred at the base of the Erythranthe group. The functionality of TR2, however, was only maintained in *M. lewisii* since it only has a sequence heterogeneous telomere while the telomeres from its sister species have only one repeat type.

Since the TR duplication occurred at the base of the Erythranthe group evolution, we investigated if TERT had undergone any evolutionary changes in the Erythranthe group. TERT orthologs were obtained from seven Erythranthe group species and from the sister clade Simiolus group we obtained the TERT orthologs from six species (Fig 1B). *M. aurantiacus* was used as the outgroup for the analysis. We first focused on the seven Erythranthe group species and conducted a molecular evolution analysis using PAML (30, 31). Results showed no evidence of rapid evolution using the branch model, site model, and a branch-site model focusing on the *M. lewisii* lineage. We then contrasted the evolutionary rate between groups using the Clade model C (CmC) (32) and tested the differences in the evolutionary rate between the Erythranthe and Simiolus group (Fig 1B). Results showed partitioning TERT into two clades fit a significantly better model compared to the null (M2a_rel_, LRT = 10.35, p-value = 5.4e-6), and the Erythranthe group had elevated levels of ⍰ (2.54) that were consistent with rapid evolution (Supplemental Table 4). We tested whether the elevated ⍰ in the Erythranthe group was due to relaxation of selection and applied the method RELAX (33). The test was not significant (K= 0.74, LRT = 0.14, p-value = 0.706) suggesting the rapid evolution of TERT was potentially due to positive selection within the Erythranthe group.

### The TRBD region of *M. lewisii* TERT can bind with both TR paralogs

Based on the molecular evolution results, we proposed the rapid evolution of TERT was driven by its molecular association with the TR duplication and predicted the *M. lewisii* TERT might have evolved unique evolutionary changes to bind with the duplicated TRs. TERT consists of four conserved domains ordered from the N- to C-terminal end as the Telomerase N-terminal (TEN) domain, Telomerase RNA-Binding Domain (TRBD), Reverse Transcriptase (RT) domain, and C-terminal Extension (CTE) domain (Fig 2A) (34). The TRBD is largely responsible for binding with the TR molecule in both animal (35–37) and plant (38) TERT. We focused on the TRBD region and tested the molecular interaction between the *Mimulus* TRBD and TR using the Yeast Three-Hybrid (Y3H) system (39). Y3H is conceptually similar to its counterpart Yeast Two Hybrid system, but it is adopted to test the interaction between a protein and RNA (Fig 2B). The system comprises of three components; first a MS2 coat protein fused with a LexA DNA-binding domain that binds with a second hybrid RNA component, which is a fusion between a MS2 stem-loop domain and a target RNA of interest. A third candidate protein fused with a Gal4 activation domain interacts with the target RNA will drive the transcription of a *HIS3* reporter gene, which can be detected by transforming an auxotrophic yeast strain (Fig 2B).

**Figure 2.**
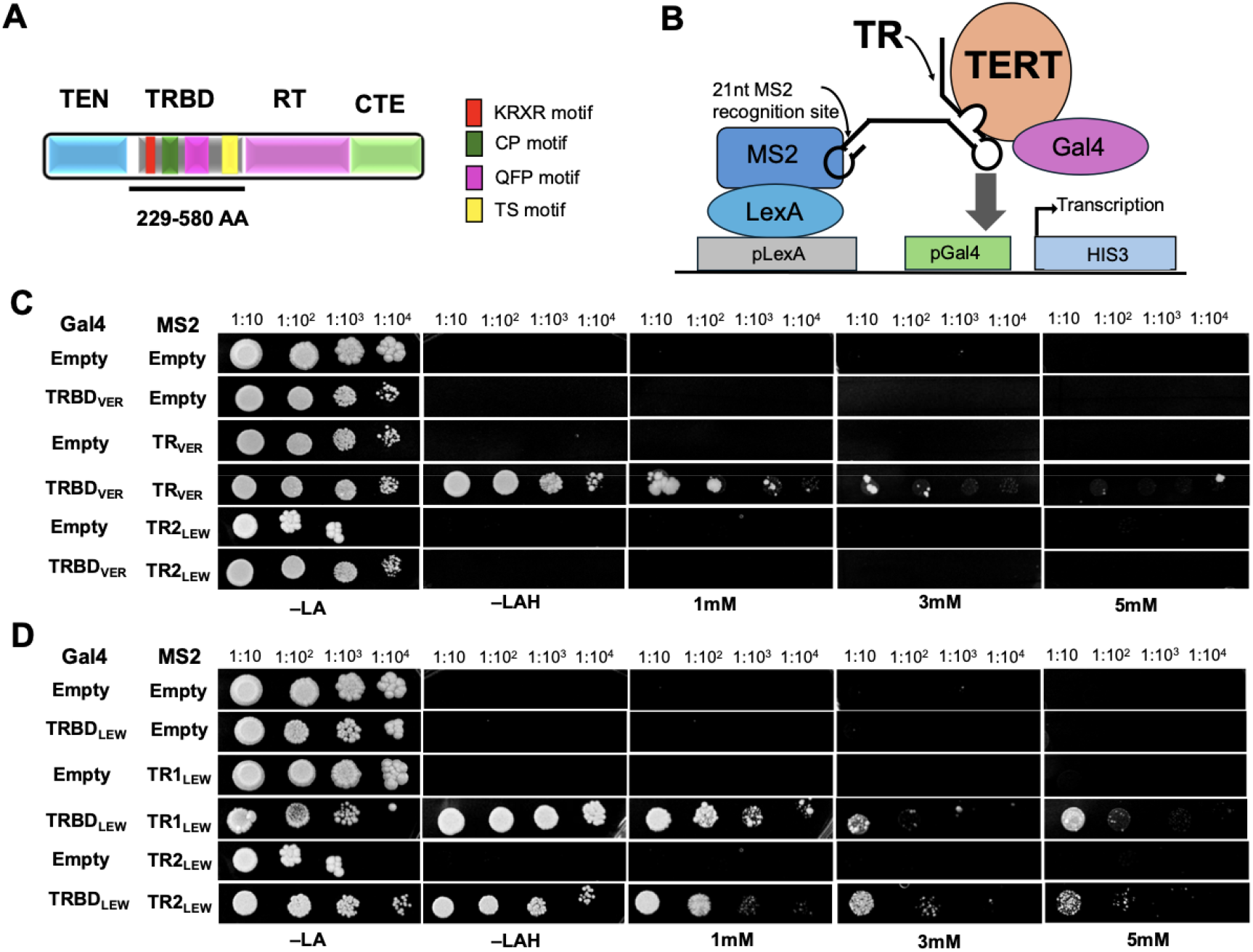
Yeast three hybrid (Y3H) testing interaction between *Mimulus* Telomerase reserve transcriptase (TERT) and telomerase RNA (TR). (A) The 4 major domains of TERT: TEN (Telomerase N-terminal; sky blue), TRBD (Telomerase RNA-Binding Domain; gray), RT (Reverse Transcriptase; pink), CTE (C-terminal Extension; green). The TRBD domain has four conserved motifs: KRxR motif, CP motif, QFP motif and TS motif. (B) The Y3H system testing interaction between the TRBD domain of TERT and TR. The MS2 homodimer protein has high affinity to a short stem-loop MS2 RNA sequence that is bound to the TR sequence. TRBD is fused to the yeast Gal4 activation domain and interaction between TRBD and TR drives the transcription of the *HIS3* reporter genes. (C) Testing the interaction of *M. verbenaceus* TRBD (TRBD_VER_) with *M. verbenaceus* TR (TR_VER_) or *M. lewisii* TR2 (TR2_LEW_). (D) Testing the interaction of *M. lewisii* TRBD (TRBD_LEW_) with TR1_LEW_ or *M. lewisii* TR2 (TR2_LEW_). Transformed yeast cells were grown on selection media and increasing concentrations of 3-amino triazole (3AT).

We first tested the binding ability of conspecific TRBD and TR combinations using Y3H (Fig 2C and D, and see Fig S1A and B for complete controls). Results showed evidence of interaction between *M. verbenaceus* TRBD (TRBD_VER_) and *M. verbenaceus* (TR_VER_) (Fig 2C lane 4), meanwhile *M. lewisii* TRBD (TRBD_LEW_) interacted with both *M. lewisii* TR1 (TR1_LEW_) and TR2 (TR2_LEW_) (Fig 2D lane 4 and 6). We repeated the experiments under increasing concentrations of 3-amino triazole (3AT), which is a chemical competitor for *HIS* biosynthesis and used for screening false positive interactions in a Y3H assay (40). All yeast transformants with the conspecific TRBD and TR combinations displayed growth under the presence of 3AT indicating the interactions were not false positive results (Fig 2C and D). In particular, the yeast transformants with the TRBD_LEW_ and TR1_LEW_ or TR2_LEW_ combinations showed similar growth under increasing 3AT concentrations (Fig 2D lane 4 and 6) suggesting TRBD_LEW_ has comparable binding strength with both TR paralogs.

The heterospecific TRBD and TR combinations were then used for testing if TRBD_VER_ can bind with TR2_LEW_, which is the recently duplicated TR paralog and functional in *M. lewisii*. Results showed no evidence of interaction between TRBD_VER_ and TR2_LEW_ (Fig 2C lane 6), suggesting *M. lewisii* TERT has potentially gained the function to bind both TR duplicates. We also tested the TRBD from *M. cardinalis* (TRBD_CAR_) since our previous study showed *M. cardinalis* had intact TR1 (TR1_CAR_) and TR2 (TR2_CAR_) paralogs, but the telomere of *M. cardinalis* consisted only of the 6bp TTTCGG repeat suggesting its TERT was not able to interact with TR2_CAR_ (22). Results showed TRBD_CAR_ can interact with TR1_CAR_ and there was growth in increasing concentrations of 3AT indicating the interaction was not false positive (Fig S1C lane 4). But for TRBD_CAR_ and TR2_CAR_ while there was growth in the -LAH selection media there was no growth under 3AT indicating TRBD_CAR_ can’t bind with TR2_CAR_ (Fig S1C lane 6). These evidences suggest *M. cardinalis* and *M. verbenaceus* TERT does not have the ability to bind with the recently derived TR2 paralog and only *M. lewisii* TERT is able to bind TR2.

### Residues near the KRxR domain underlie the dual TR binding

We narrowed down the TRBD_LEW_ region that was responsible for binding TR2_LEW_ using a domain swap experiment. Two domain swap TRBD constructs were designed (Fig 3A). First construct consisted of a fusion between the N-terminal TRBD_LEW_ with the C-terminal TRBD_VER_ (TRBD_LxV_), and the second construct consisted of a fusion between the N-terminal TRBD_VER_ with the C-terminal TRBD_LEW_ (TRBD_VxL_). The C-terminal end of the TRBD holds the CP, QFP, and T motifs that are conserved across eukaryotes (41), while the N-terminal side holds a KRxR motif recently discovered in *A. thaliana* (38). Both TRBD constructs were tested with TR2_LEW_ (Fig 3B and see Fig S2 for complete controls). Results showed TRBD_LxV_ was able to interact with TR2_LEW_ and with increasing concentration of 3AT there was still yeast growth (Fig 3B lane 4). Noticeably there was growth at 5mM 3AT which was also observed for the Y3H assay testing the interaction between TRBD_LEW_ and TR2_LEW_ (Fig 2D lane 6). On the other hand, TRBD_VxL_ showed very weak or no interaction with TR2_LEW_ (Fig 3B lane 6), suggesting the N-terminal region of *M. lewisii* TRBD contributes to binding with TR2.

**Figure 3.**
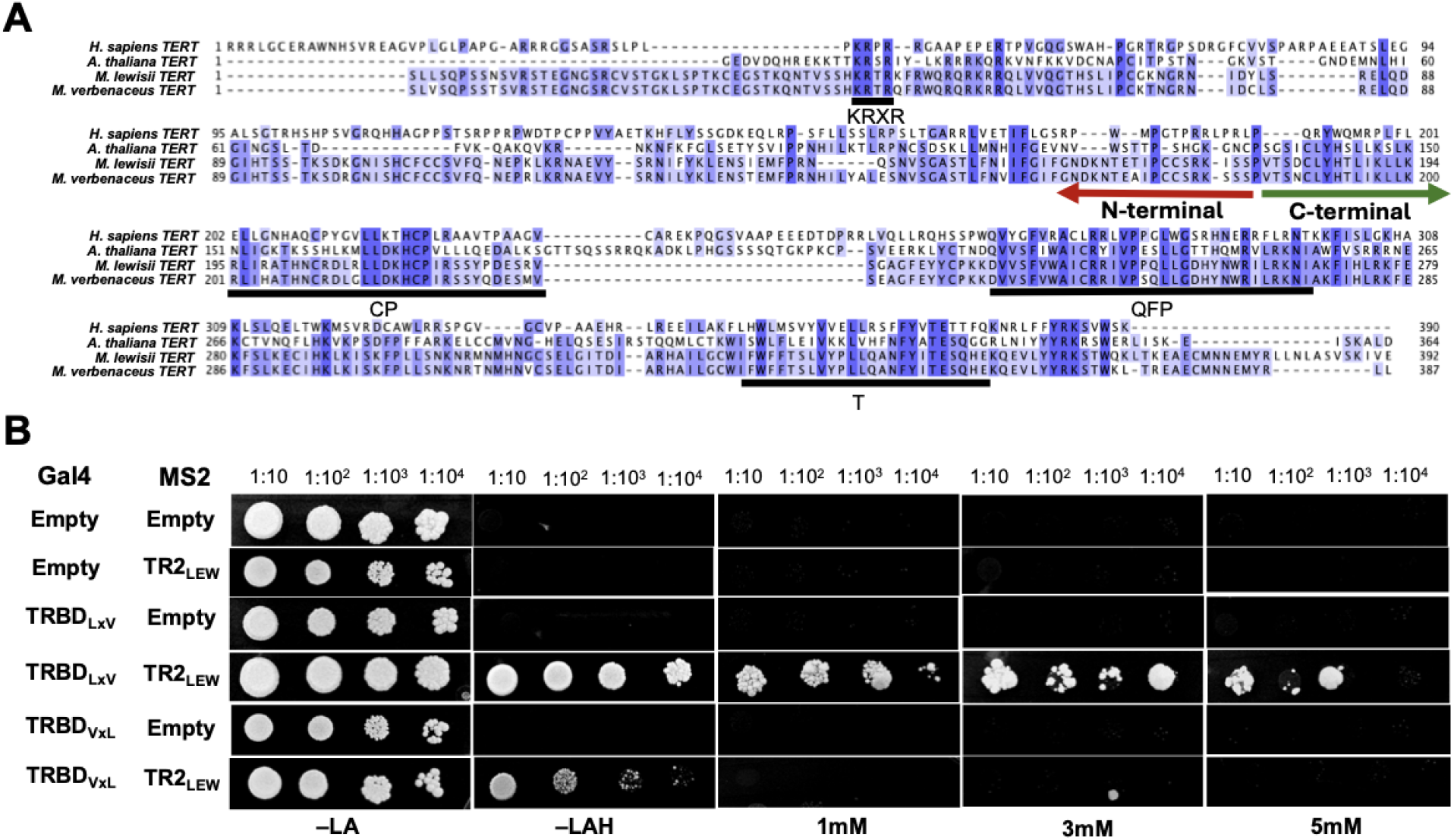
The N-terminal region of *M. lewisii* TRBD binds with the duplicated TRs. (A) Multisequence alignment of TRBD from human, *A. thaliana*, *M. lewisii,* and *M. verbenaceus*. Arrows indicate the demarcation of the N– and C–terminal region of the TRBD used for the domain swap experiment. (B) Yeast three-hybrid experiment testing the binding between domain swapped TRBD and *M. lewisii* TR2 (TR2_LEW_). Transformed yeast cells were grown in selection plates and increasing concentration of 3-amino triazole (3AT).

To identify the amino acids within the N-terminal region that were responsible for TR binding, we first compared the TRBD sequences across multiple *Mimulus* species to identify the candidate residues (Fig 4A). At position 277 *M. lewisii* had a lysine (K) whereas all other Erythranthe group species had a glutamine (Q), and at position 309 *M. lewisii* had a tyrosine (Y) whereas all other Erthranthe group species had a cysteine (C). We tested the functions of these substitutions by creating three constructs that mutated TRBD_LEW_. The first construct changed position 277 K to Q (TRBD_K277Q_), the second construct changed position 309 Y to C (TRBD_Y309C_), and the third construct changed both amino acids (TRBD_K277Q+Y309C_). These constructs were tested against TR1_LEW_ and TR2_LEW_ using Y3H (Fig 4B and see Fig S3 for complete controls). Results showed no evidence of interaction for TRBD_K277Q+Y309C_ (Fig 4B lane 11 and 12) and when we tested the single amino acid substitutions separately, TRBD_K277Q_ showed interaction in the selection media (–LAH) but neither TRBD_K277Q_ (Fig 4B lane 5 and 6) or TRBD_Y309C_ (Fig 4B lane 8 and 9) showed interaction in the presence of 3AT. These results suggest K277 and Y309 amino acids are crucial for binding both TR paralogs for the *M. lewisii* TERT.

**Figure 4.**
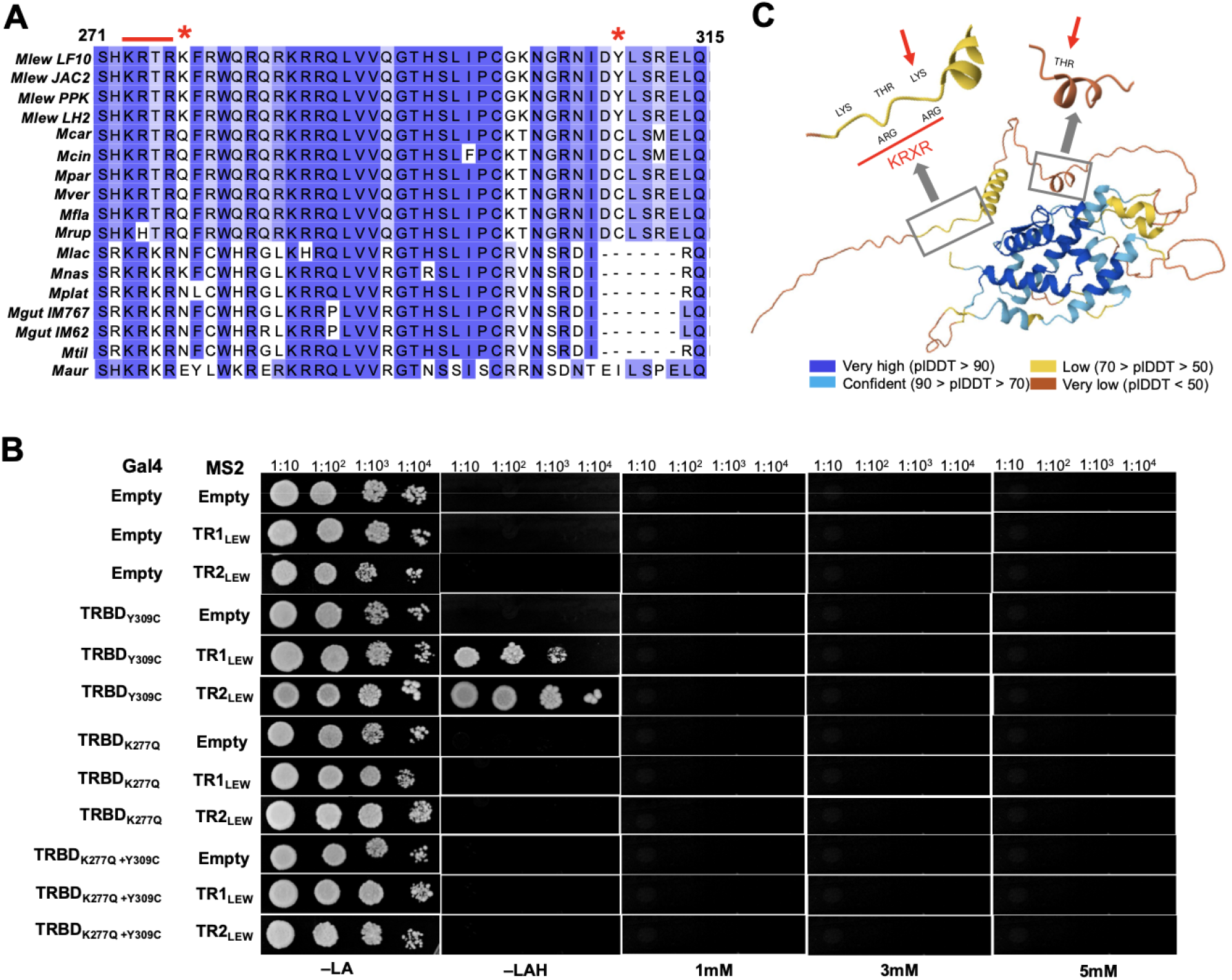
Two amino acids in *M. lewisii* are associated with the dual TR binding. (A) Multisequence alignment of the KRxR motif (red line) region in *Mimulus*. Shown are sequences from *M. lewisii* (Mlew) genotypes LF10, JAC2, PPK, and LH2. Additional sequences from species *M. cardinalis* (Mcar), *M. cinnabar* (Mcin), *M. parishii* (Mpar), *M. verbenaceus* (Mver), *M. flammea* (Mfla), *M. rupestris* (Mrup), *M. laciniatus* (Mlac), *M. nasutus* (Mnas), *M. platycalyx* (Mpla), *M. guttatus* (Mgut) genotypes IM767 and IM62, *M. tilingii* (Mtil), and *M. aurantiacus* (Maur). The two amino acids of interest are indicated with red stars. (B) Yeast three-hybrid experiments testing TRBD with substitutions of the two amino acids and *M. lewisii* TR1 or TR2. (C) AlphaFold model of *M. lewisii* TRBD. The KRxR motif is shown with red underline. The red arrowheads indicate the lysine (K) position 277 and tyrosine (Y) position 309. Confidence of the protein folding is indicated with the plDDT scores.

AlphaFold was used to predict the structural model of the *M. lewisii* TRBD and explore the three dimensional structure involving the K277 and Y309 amino acids (Fig 4C). Results showed the C-terminal end was well folded with high pLDDT scores (plDDT > 70) but the N-terminal end had low confidence scores and was largely unstructured. This region is known to have poor conservation across eukaryotic TERT sequences and contains largely lineage specific motifs (42). The K277 and Y309 amino acids flank the KRxR motif, suggesting amino acid changes in the evolutionarily flexible region of TRBD may underlie the TR binding flexibility of the *M. lewisii* TERT.

### Quantifying the telomerase activity by combining TRAP with next generation sequencing

We next investigated if the telomerase of *M. lewisii* had evolved a different activity compared to species without the TR duplication using TRAP. The activity of the telomerase can be divided into two categories (43); processivity measures the number of telomeric repeats synthesized onto the substrate and productivity measures the number of substrate molecules extended by the telomerase. Previously, we conducted TRAP on *M. lewisii* and showed its telomerase synthesized both 6 bp TTTCGG and 7bp TTTCGGG repeats (22). But because we used Sanger sequencing on a select number of TRAP products our conclusions were incomplete. To characterize the telomerase processivity and productivity of *M. lewisii* and its sister species, we combined TRAP with next generation sequencing (TRAP_NGS_) and bioinformatically inferred the activity of the telomerase.

After TRAP the concentration of the resulting products is often low and requires PCR to amplify the products for downstream analysis (*e.g.* gel electrophoresis or quantitative PCR). To investigate any potential PCR amplification bias we performed TRAP_NGS_ on *M. lewisii* TRAP products after 5, 10, 15, and 20 PCR cycles. Results showed the increasing PCR cycles led to an exponential increase in TRAP sequences, where there was a 1000 fold difference in TRAP products between 5 and 20 PCR cycles (Fig S4A). We then assumed the *M. lewisii* telomerase bound with a TR1 paralog synthesized a product with the 6bp TTTCGG repeat, while a telomerase bound with a TR2 paralog synthesized a product with the 7bp TTTCGGG repeat. Each sequencing read was classified either as a 6bp TTTCGG or 7bp TTTCGGG repeat product. We investigated the number of repeats for each TRAP product and noticed at 5 PCR cycles no 7bp TTTCGGG TRAP products were detected (Fig S4B), likely because its products are less in quantity compared to the 6 bp TTCGGG TRAP products (22). With increasing PCR cycle the mean number of repeats were decreasing, suggesting the PCR step was amplifying lower molecular size DNA products. We chose 15 PCR cycles for downstream TRAP_NGS_ experiments because it provided sufficient amplification of TRAP products while preserving the relative differences between 6 bp TTTCGG and 7bp TTTCGGG telomerase extension products (Fig S4B).

We conducted TRAP_NGS_ on *M. lewisii* and compared the results to its sister species *M. cardinalis* and *M. verbenaceus* which does not have a functioning TR2 paralog. TRAP products of the three species were initially visualized on a polyacrylamide gel that showed a ladder of bands corresponding to the telomerase products differing in size due to different numbers of telomere repeats (Fig S5A). We noticed the intensity of the bands differed between species, where *M. lewisii* had the strongest intensity. After applying TRAP_NGS_ on the telomerase products we bioinformatically inferred the processivity and productivity from the resulting sequencing reads (Fig 5).

**Figure 5.**
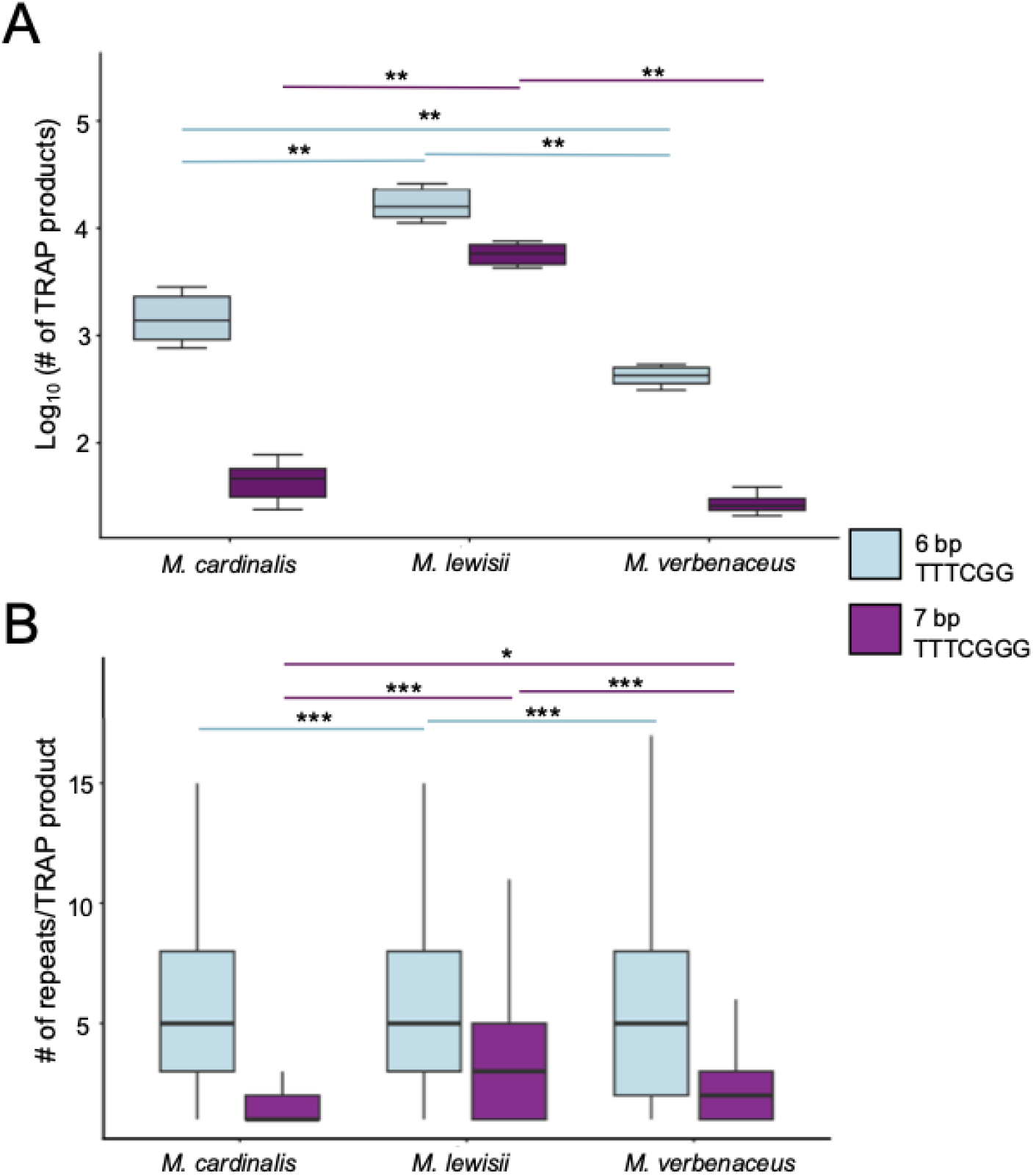
TRAP_NGS_ shows elevated telomerase activity for *M. lewisii*. (A) Number of TRAP products that were categorized as a product of TR1 and contains 6bp TTTCGG repeats (blue) or categorized as a product of TR2 and contains 7bp TTTCGGG repeats (purple). (B) Number of repeats per TRAP product. *,**,*** indicate significant differences p < 0.05, < 0.01, < 0.001 respectively after Mann–Whitney U test.

For the 6bp TTTCGG repeat products, *M. lewisii* had a significantly higher number of products (Mann-Whitney U test p-value < 0.01) compared to *M. cardinalis* and *M. verbenaceus* (Fig 5A). For the 7bp TTTCGGG repeat products, *M lewisii* also had a significantly higher number of products (Mann-Whitney U test p-value < 0.01). The amount of 7bp TTTCGGG repeat products were very low in *M. cardinalis* and *M. verbenaceus* (100 fold less than their 6bp TTTCGG TRAP products). Because *M. cardinalis* and *M. verbenaceus* were not predicted to synthesize the 7bp TTTCGGG repeats, the low abundance of these repeats suggested these products are either errors of the telomerase or the TRAP_NGS_ assay. Overall, the total number of TRAP products was significantly higher in *M. lewisii* (Mann-Whitney U test p-value < 0.001) and greater than 10-fold compared to *M. cardinalis* and *M. verbenaceus* (Fig S5B). Within *M lewisii* the 6bp TTTCGG repeat had a significantly higher number of TRAP products compared to the 7bp TTTCGGG repeat (Mann-Whitney U test p-value < 0.01), suggesting a difference in productivity between the TR1 and TR2 paralogs by the *M. lewisii* telomerase.

To validate the elevated telomerase productivity we repeated the experiment using an internal *E. coli* genomic DNA spike-in as a normalization factor between libraries. Consistent with the initial TRAP_NGS_ experiments, *M. lewisii* showed significantly higher levels of normalized telomerase productivity compared with *M. cardinalis* and *M. verbenaceus* (Fig S5C Mann–Whitney U test, p < 0.01). *M. lewisii* produced approximately 13-fold and 59-fold more telomerase products than *M. cardinalis* and *M. verbenaceus*, respectively. These results confirm the elevated TRAP_NGS_ signal in *M. lewisii* reflects increased telomerase productivity rather than differences in sequencing depth or technical variation during library preparation. The spike-in normalized data also revealed both telomerase products generated by *M. lewisii* were elevated relative to the other species (Fig S5D). TR1 derived products were approximately 10–50-fold higher in *M. lewisii* compared with *M. cardinalis* and *M. verbenaceus*, while TR2 derived products were detected at substantially higher levels only in *M. lewisii* (Mann–Whitney U test, p < 0.001). Also consistent with the initial TRAP_NGS_ results, within *M. lewisii* the TR1 derived products were significantly more abundant than TR2 derived products (Mann–Whitney U test, p < 0.01).

We next quantified the number of telomere repeats for each TRAP product to estimate the processivity of the telomerase for each species (Fig 5B). We sequenced three biological replicates for each species, and the replicates showed a consistent number of repeat counts (Fig S5E). For the 6bp TTTCGG repeat products, the mean repeat number per TRAP product were similar among the three species (*M. lewisii* = 5.7, *M. cardinalis* = 5.5, and *M. verbenaceus* = 5.5) and it was significantly elevated for *M. lewisii* (Mann-Whitney U test p-value < 0.001). For the 7bp TTTCGGG repeat products, *M. lewisii* had almost twice the number of mean repeats per TRAP product (*M. lewisii* = 3.6, *M. cardinalis* = 1.8, and *M. verbenaceus* = 2.0), further suggesting the 7bp TTTCGGG repeat products for *M. cardinalis* and *M. verbenaceus* were likely minor products of their telomerase. Within *M. lewisii* the 6bp TTTCGG repeat products had significantly higher number of repeats per TRAP product (Mann-Whitney U test p-value < 0.001) compared to the 7bp TTTCGGG repeat products. This suggested *M. lewisii* telomerase has a higher processivity using the TR1 paralog compared to the TR2 paralog.

### Transcriptomic analysis reveals limited expression divergence of telomere maintenance genes in *M. lewisii*

The TRAP_NGS_ analysis revealed an enhanced telomerase activity in *M. lewisii.* We investigated if this elevated activity might be associated with a divergence of gene expression in *M. lewisii*, by performing a comparative transcriptomic analysis of the apical meristem (AM) and seedling tissues from *M. cardinalis*, *M. lewisii*, *M. parishii*, and *M. verbenaceus*. We focused on these four species to contrast *M. lewisii* to its sister species without a functioning TR2 duplication, and we contrasted the transcriptome of the seedling to the AM since the telomerase is most active in the meristem (Fig 5 and (22)). Transcriptomes for each species were aligned to its own reference genome and we were able to align over 96% of the total reads, except for 1 library from *M. verbenaceus* with 83% alignment percentage. (Supplemental Table 5).

We estimated gene expression values for 16,853 orthogroups representing 1 to 1 orthology among all four *Mimulus* species. An initial principal component analysis (PCA) of all transcriptomes revealed clear separation of samples according to tissue type and species identity (Fig S6). PC1 primarily separated AM and seedling tissues and PC2 captured variation among species. Biological replicates clustered closely within their corresponding species–tissue groups, demonstrating strong reproducibility among samples. We then focused our analysis on the expression pattern of telomere maintenance genes. Initially we examined if telomere maintenance genes were highly expressed in the AM compared to seedling by conducting a differential gene expression analysis (Fig 6A). Within species, the majority of the telomere maintenance genes had significant differential expression in the AM compared to seedling tissue (13 genes in *M. cardinalis*, 9 genes in *M. lewisii*, 13 genes in *M. parishii*, and 12 genes in *M. verbenaceus*). In all four species the genes TERT, NBS1, KU80, POT1B, MRE11, and CTC1 showed a significant upregulation in the AM, indicating enhanced expression of telomere maintenance genes are a conserved feature of AM tissue.

**Figure 6.**
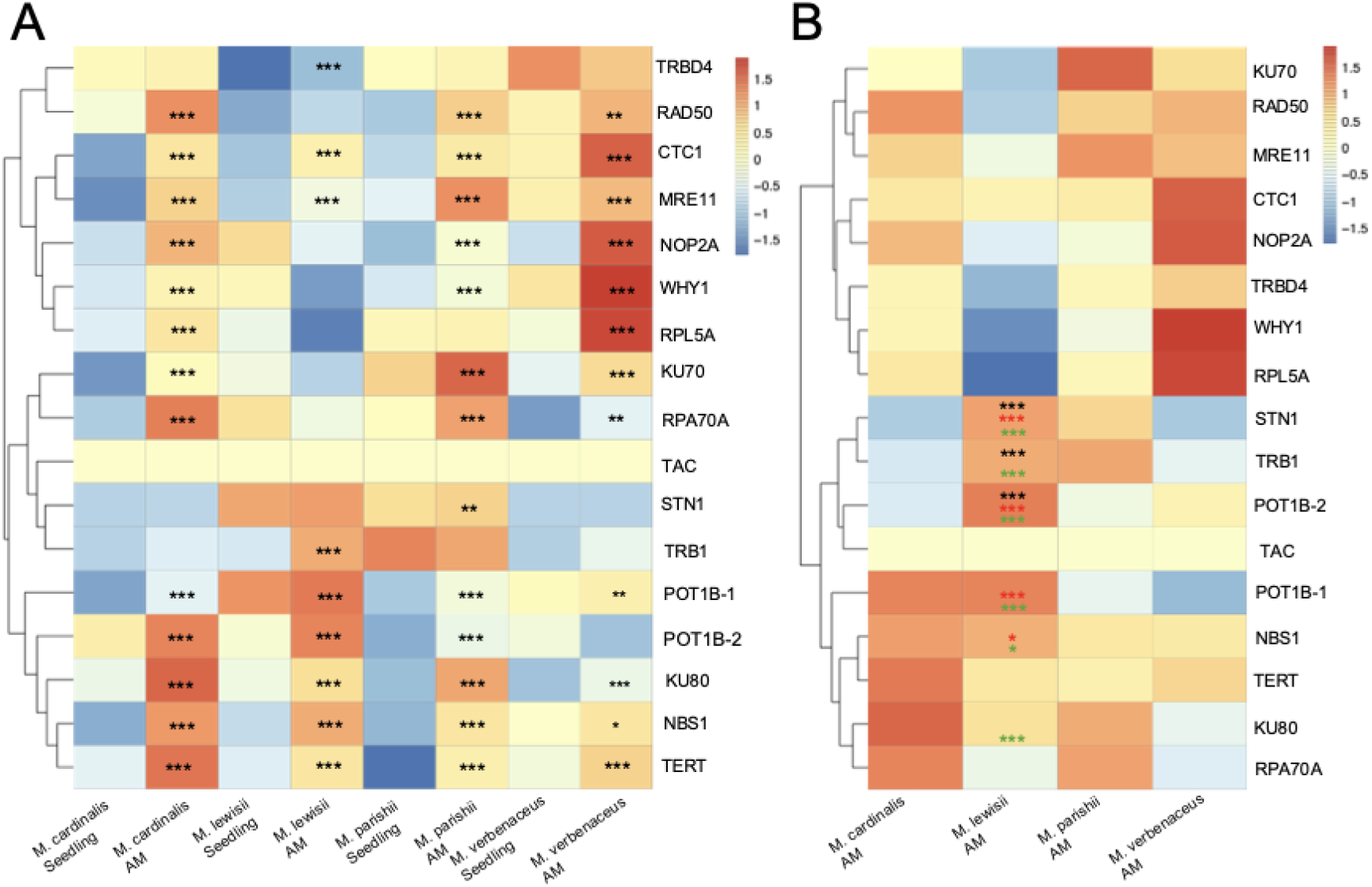
Gene expression of telomere maintenance genes in *M. cardinalis*, *M. lewisii*, *M. parishii*, and *M. verbenaceus*. Shown are heatmap of log CPM values averaged across biological replicates and after z-score transformation. A) Within species differential expression of telomere maintenance genes between the apical meristem (AM) and seedling. Genes with significant differential gene expression in the AM are labeled with a star (*). B) Between species differential expression of telomere maintenance genes in the apical meristem. Black, red, and green star (*) indicate genes with significantly higher expression in *M. lewisii* relative to *M. cardinalis*, *M. verbenaceus*, or *M. parishii* respectively. *,**,*** indicate significant differences p < 0.05, < 0.01, < 0.001 after Benjamini–Hochberg false discovery rate.

We then compared the expression of telomere maintenance genes in the AM between species. Specifically we asked if telomere maintenance genes in *M. lewisii* had higher expression in the AM compared to other *Mimulus* species. Pairwise comparison against *M. lewisii* AM revealed only three telomere maintenance genes (STN1, TRB1, and POT1B) had significantly higher expression in *M. lewisii* compared to all three other *Mimulus* species (Fig 6B). Importantly TERT, which is the main catalytic protein of the telomerase, was not significantly different in its expression. These results indicate that elevated telomerase activity in *M. lewisii* is not explained by broad transcriptional upregulation of telomere maintenance genes.

### Telomere sequence analysis in F1 hybrids between *M. lewisii* and *M. verbenaceus*

Our results suggest *M. lewisii* TERT has gained functions to interact with the duplicated TRs and has increased telomerase activity compared to its sister species without the TR duplication. We further investigated the *in vivo* functionality of the *M. lewisii* telomerase by studying the telomeres of a F1 hybrid generated from crossing *M. lewisii* with *M. verbenaceus*, which does not have the duplicated TR2 paralog and synthesizes a telomere consisting only of the 6bp TTTCGG repeat. During the telomere maintenance of the F1 hybrid, we hypothesized the telomerase of the *M. lewisii* and *M. verbenaceus* would genetically interact with each other resulting in two potential outcomes (Fig 7A). In the telomere persistence model, we propose a species specific telomerase and telomere DNA interaction maintains the F1 chromosome ends resulting in the telomere sequence unchanged from the parental telomeres. Meanwhile in the telomere conversion model, we propose a difference in telomerase activity between *M. lewisii* and *M. verbenaceus*, and the F1 chromosome ends will convert into the parental telomere with the dominant (44) telomerase. We investigated the two models by nanopore sequencing the F1 progeny and classifying the telomere reads by its originating species specific chromosome ends.

We nanopore sequenced a F1 hybrid plant generating 6.1 Gbp of sequencing data with read length N50 of 13.4 kbp (Supplemental Table 6). The data were comparable to the nanopore long read data of its parents (Supplemental Table 6). The long reads were then aligned to the *M. lewisii* or *M. verbenaceus* reference genome to extract species specific telomere long reads. There were 52 reads uniquely aligned to *M. lewisii* chromosome ends and 61 reads uniquely aligned to *M. verbenaceus* chromosome ends. The telomere long reads from the F1 were then compared to the nanopore sequencing data from the parents. We focused on the 7 bp TTTCGGG repeat since it can only be synthesized by the *M. lewisii* telomerase, its quantity can be used to approximate the activity of the *M. lewisii* telomerase. Initially we compared the telomere long reads of the F1 hybrid originating from the *M. lewisii* chromosomes (F1_LEW_) and the telomere long reads from the *M. lewisii* parent (P_LEW_). Results showed both F1_LEW_ and P_LEW_ telomere long read sequences consisted of a mix of 6 bp TTTCGG and 7 bp TTTCGGG repeats (Fig 7B and C) and there was no significant difference in the amount of 7 bp TTTCGGG repeat (Fig S7A). Comparing telomere long reads of the F1 hybrid originating from the *M. verbenaceus* chromosomes (F1_VER_) and telomere long reads from the *M. verbenaceus* parent (P_VER_), the F1_VER_ telomere long reads had noticeable amounts of 7 bp TTTCGGG repeats (Fig 7B and C) and this was significantly higher compared to P_VER_ telomere long reads (Fig S7A, Mann-Whitney U test p-value < 0.001). But when the amount of 7 bp TTTCGGG repeats were compared to F1_LEW_ and P_LEW_ telomere long reads, F1_VER_ had significantly lower amounts (Fig S7A, Mann-Whitney U test p-value < 0.001).

**Figure 7.**
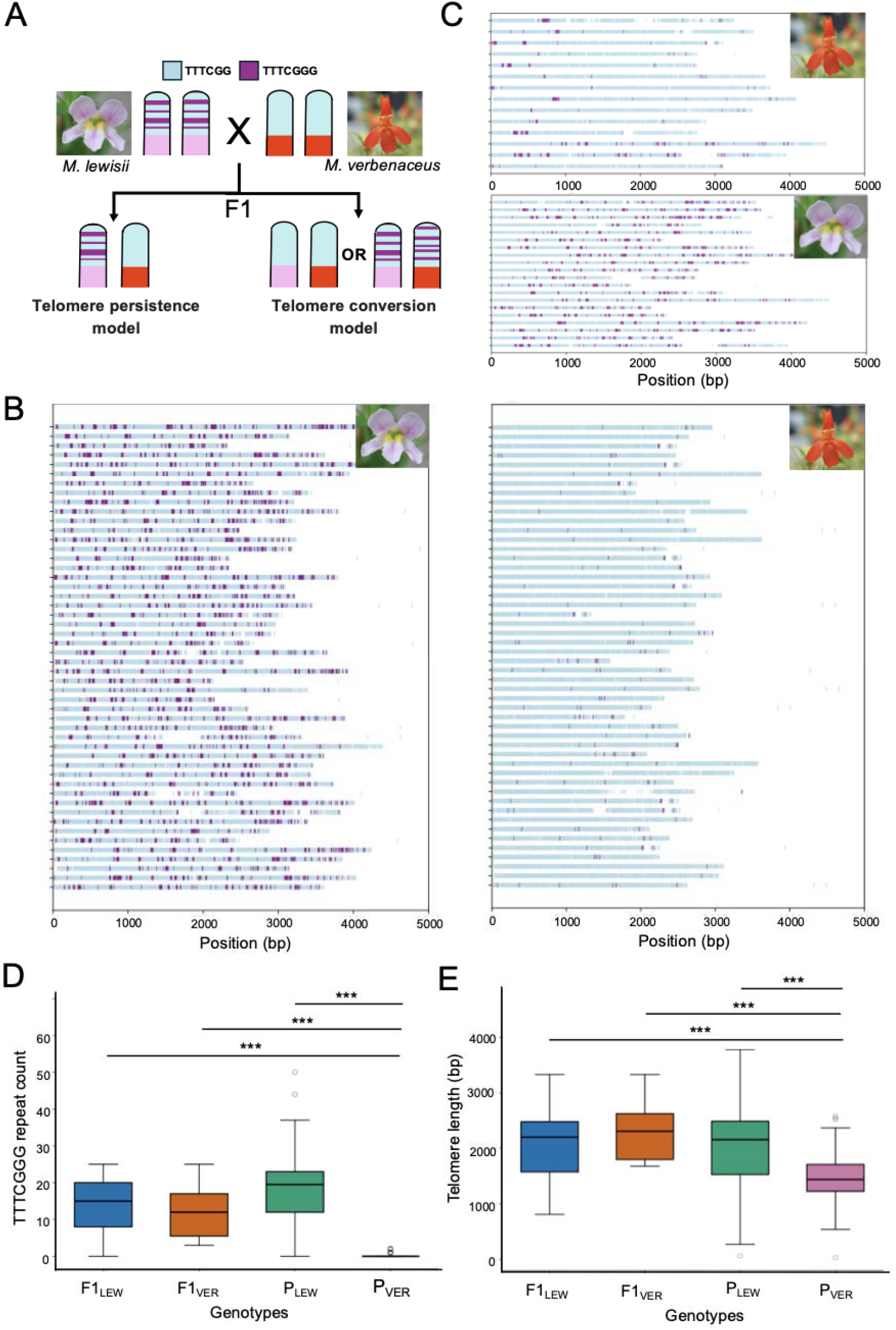
Quantifying telomere repeat patterns and telomere length estimates in the F1 interspecies hybrid. A) Two possible consequences in the telomeres of the F1 hybrid crossed between *M. lewisii* and *M. verbenaceus*. B) Telomeric repeat patterns from long reads sequenced and originating from the telomeres in the parental *M. lewisii* plant (left) and *M. verbenaceus* plant (right). C) Telomeric repeat patterns from long reads sequenced from a F1 plant. Shown are telomere long reads originating from the *M. lewisii* chromosomes (bottom) and *M. verbenaceus* chromosomes (top). D) Boxplot quantifying the amount of 7bp TTTCGGG repeat within the first 1kbp of a telomere long read from *M. lewisii* chromosomes of the F1 (F1_LEW_) and parent (P_LEW_), and the *M. verbenaceus* chromosomes of the F1 (F1_VER_) and parent (P_VER_). E) Boxplot shows the telomere length estimates from the telomere long reads originating from the F1_LEW_, P_LEW_, F1_VER_ and P_VER_ chromosomes. *** indicate significant differences (p < 0.001) after Mann–Whitney U test.

We hypothesized the increased 7 bp TTTCGGG repeats in the F1_VER_ telomere long reads were newly synthesized telomere DNA in the F1 hybrid. In a TERT null *A. thaliana* mutant, its telomere loses 500 bp of DNA every generation (45), suggesting a couple of hundreds of basepairs are elongated in plants every generation. We focused our analysis of the 7 bp TTTCGGG repeat counts in the first 1 kbp of each telomere long read as this region would contain newly synthesized telomere DNA repeats. Comparing the telomere long reads in the F1 hybrid, results showed no significant differences in the 7 bp TTTCGGG repeats in the first 1 kbp of F1_LEW_ and F1_VER_ telomere long reads, but both had significantly higher 7 bp TTTCGGG repeats compared to P_VER_ telomere long reads (Fig 7D, Mann-Whitney U test p-value < 0.001). This suggested telomere maintenance in the F1 hybrid was most consistent with the telomere conversion model (Fig 7A). We further investigated the telomere conversion model by examining the telomere length of the F1 hybrid. Previously, we discovered *M. lewisii* had a 20% longer telomeres compared to *M. verbenaceus* (22), and we asked if the telomeres in the F1_VER_ chromosome ends had elongated as well. Results showed telomeres from P_VER_ were significantly shorter compared to telomeres from F1_LEW_, P_LEW_, and F1_VER_ (Fig 7E Mann-Whitney U test p-value < 0.001), indicating at the F1 chromosome ends the telomeres of *M. verbenaceus* chromosomes had converted into the telomeres of *M. lewisii* chromosomes.

## Discussion

In this study we have investigated the molecular evolution of the *Mimulus* TERT and the functional interaction it has with TR. The TR gene is known to evolve rapidly at the sequence level while retaining conserved structural elements (46, 47). In *M. lewisii* the TR gene had duplicated providing a unique opportunity to investigate how duplication and sequence divergence at the RNA component of the telomerase influenced the evolution of its catalytic protein partner TERT. Our results showed the TR duplication occurred deep in the ancestral Erythranthe common ancestor and TERT is rapidly evolving across the Erythranthe group (Fig 1B), suggesting the duplication of the TR drives the coevolution of TERT. However, only in *M. lewisii* the duplicated TR paralogs were able to interact with TERT (Fig 2D lane 4 and 6), raising the question why does TERT show elevated evolutionary rates across the entire Erythranthe lineage? We suggest the rapid TERT evolution across the lineage may represent a broader history of adaptation to a dynamic TR evolution, with *M. lewisii* retaining a unique example where this evolutionary flexibility is revealed through the coexistence of two sequence divergent TR paralogs.

Example cases of RNA binding proteins evolving to expand its repertoire of RNA are seen between the bacterial coat protein PP7 and RNA bacteriophages (48), La-related proteins and transcripts from RNA polymerase III (49), and the argonaute protein and small RNAs (50). One way these RNA binding proteins are able to bind with a diversity of RNA species is by recognizing the secondary structure of the RNA (51, 52). The eukaryote TR contains two core structural domains: a template-proximal pseudoknot (PK) structure and a template-distal helical domain P4/5/6 that is crucial for telomerase activity (53). Previously, yeast three-hybrid assays testing the binding activity of *A. thaliana* TERT-TR showed the conserved TRBD region interacted with the PK structure while the KRxR motif interacted with P4 stem structure (38). The KRxR motif is not only conserved between *Mimulus* and *Arabidopsis* TERT but also between *Mimulus* and human TERT as well (Fig 3A). We found *M. lewisii* has species specific lysine and tyrosine amino acids flanking the KRxR motif and substituting these to glutamine and cysteine respectively ablated its TR binding ability. Lysine is a predominantly favored amino acid in protein-RNA interactions (54) while tyrosine has a preference for binding with adenosine nucleotides (55). Substitutions involving these amino acids near the KRxR motif might explain the ability of the *M. lewisii* TERT to bind two sequence divergent RNA molecules. We note TERT is rapidly evolving in the entire Erythranthe group, and we hypothesize in lineages without a functioning TR2 the rapid evolution is driven to avoid the interaction with the duplicated TR2. The molecular evolution of TERT may therefore reflect adaptation to maintain functional compatibility with a rapidly evolving TR landscape, mainly to gain or lose the interaction with the derived TR duplicate. What selective pressure is driving the rapid TERT-TR coevolution in *Mimulus* remains to be known.

The functional consequence arising from duplicated TR is not known in plants, but hints can be gained from mammalian studies. Plants share a telomerase-based mechanism of telomere regulation with the majority of animals, including mammals(56). In normal mammalian cells, increased cellular replication results in a gradual shortening of the telomere until it reaches a catastrophic length that triggers genomic instability and ultimately apoptosis (*i.e.* replicative senescence) (57). Tumor cells, on the other hand, suppress cellular senescence and death, and one way to achieve this is through increasing the activity of the telomerase and maintaining the telomere from shortening (58). A variety of human cancer cells upregulates the expression of the TR gene (59–62) and in some cases even amplifies the TR locus (63). In plants, tumors are relatively rare and not as lethal as animal tumors (64), indicating an increased telomerase activity in plant cells will not have the same deleterious effects as in animals. We propose the TR duplication in *M. lewisii* is functionally linked to the telomerase activity. Our TRAP_NGS_ results suggested the telomerase of *M. lewisii* had over 10 fold levels of productivity compared to its sister species *M. cardinalis* and *M. verbenaceus* that do not have a functioning TR2. Similar results are seen in Marek’s Disease Virus where an integration of a viral TR from a herpes virus can increase the telomerase activity in chicken (65, 66). But what consequence does the higher telomerase activity have on the plant? Overexpressing TERT and TR in *A. thaliana* can significantly increase the activity of the telomerase, but no corresponding increase in telomere length has been observed (67). In *M. lewisii,* we found no significant evidence of expression divergence of TERT compared to its sister species (Fig 6B), meanwhile *M. lewisii* had significantly higher telomerase activity and longer telomeres than its sister species without a functioning TR2 (Fig 7B). This suggests unlike *A. thaliana*, the activity of the telomerase could be linked to telomere length regulation in *M. lewisii*. Future study would be needed to test the relationship between TR duplication and telomere length maintenance.

Length of the telomere varies naturally between individuals (53, 68) and what determines this variation has been a largely open question. In *A. thaliana*, telomere length has a high broad-sense heritability indicating genetic variation between individuals can have a large effect on the length variation (69). But telomeres are also directly inherited from the parents and it’s unclear how much change could happen in the telomere between the parent and the offspring. For example in *A. thaliana*, a F1 cross between a long and short telomere parent, the telomere length of the hybrid is bimodal suggesting direct inheritance of the telomere with minimal change in its length (70). But in mice, offspring telomere length is determined through a parent-of-origin effect, where it can be elongated if the maternal telomere length was short and the paternal telomere length was long (71). Here, we suggest another model of telomere length inheritance consistent with a dominance effect (44, 72). *M. lewisii* telomerase is dominant over the *M. verbenacus* telomerase during the maintenance of the F1 offspring telomere, and the ends of the *M. verbenacus* chromosome had phenotypically converted into telomeres of *M. lewisii*. Interchromosomal dominance effects are also observed in the nucleolus where the rDNA cluster of one chromosome becomes silenced in genetic hybrids (73). While nucleolar dominance is mediated through the epigenetic pathway (74), the telomerase dominance effect we observe is a trans effect of the telomerase complex. The ability to overwrite and change the composition of the telomere in one generation suggests the telomere composition of *M. lewisii* could easily spread across the Erythranthe group. Genome-wide analysis shows extensive evidence of hybridization that has occurred between *M. lewisii* and other species of the Erythranthe group (75). But despite this hybridization only *M. lewisii* has maintained the TR duplication and sequence heterogeneous telomere, suggesting the *M. lewisii* telomeres might be selected against in other Erythranthe group species. Elucidating the evolutionary pressure responsible for the dichotomous telomere sequence structure in the Erythranthe group will be an important direction for future research.

## Materials and method

### Nanopore sequencing Erythranthe groups species

High molecular weight (HMW) DNA was extracted for each plant from leaf tissues using the Wizard HMW DNA Extraction Kit (Promega catalog #A2920) according to the manufacturer’s instructions. Nanopore sequencing library were prepared following the Ligation Sequencing Kit V14 (Oxford Nanopore Technologies catalog #SQK-LSK114). For each library, 1 μg of genomic DNA was used as input for DNA repair and end preparation. DNA repair was performed using the NEBNext FFPE DNA Repair Mix (New England Biolabs catalog #M6630) and the Ultra II End Prep Enzyme Mix (New England Biolabs catalog #E7546), followed by purification using AMPure XP Beads and elution in 61 μL of nuclease-free water. Adapter ligation was performed by combining 60 μL of purified DNA with 5 μL of ligation adapter using the NEBNext Quick T4 DNA Ligase (New England Biolabs catalog #E6056) for 10 min at room temperature. Ligated DNA was purified using AMPure XP beads and washed twice with Long Fragment Buffer. Final libraries were eluted in 15 μL of elution buffer and quantified using a Qubit fluorometer prior to sequencing.

Sequencing was performed on FLO-MIN114 flowcells with R10.4.1 nanopores and sequenced using the MinION Mk1D sequencer. A total of 1000 μL of flow cell priming mix was loaded through the priming port, and 75 μL of library mixed with Library Beads was loaded onto the SpotON sample port for sequencing. After sequencing was completed basecalling on the raw nanopore sequencing POD5 files were carried out with dorado v0.9.6 (https://github.com/nanoporetech/dorado) using duplex mode and super-accurate (sup) model.

### Nanopore long read sequencing analysis of telomere repeats across genotypes and species

Using the newly sequenced nanopore sequencing data the telomere repeats were quantified across genotypes and species using the raw sequencing long reads. We used the Topsicle package (29) to find long reads originating from the telomere region using a TRC threshold of 0.7. For each telomere long read the counts for the 6bp TTTCGG repeat were made by matching the sequence TTTCGGT, while the counts for the 7 bp TTTCGGG repeat were made by matching the sequence TTTCGGG. The proportions of 6bp and 7bp of each species were calculated by dividing the number of matches in each category to the total matches.

### Generating 17-way whole-genome alignments

Raw nanopore sequencing reads were initially filtered to select for reads that were longer than 10kbp and have an average q-score greater than 9 using the chopper program (76). Filtered reads were used for *de novo* genome assembly using Flye (v2.9.5) (77) and polished with medaka (v2.1.1) (https://github.com/nanoporetech/medaka). Genome assemblies for *M. aurantiacus* (78), *M. cardinalis*, *M. lewisii*, *M. parishii*, and *M. verbenaceus* were downloaded from Mimubase (http://mimubase.org/). Genome sequences for *M. laciniata* were obtained from the Sequence Read Archive with identifier PRJNA1060722 (79). For each genome assembly repetitive DNA were identified using RepeatModeler (v2.0.7) (80) and softmasked using RepeatMasker (v4.2.1) (http://www.repeatmasker.org/). To generate an initial guide tree for genome alignments mashtree (v1.4.6) (81) was used and rooted based on *M. aurantiacus*. The softmaked genome assemblies were used to construct a multi-genome alignment using Progressive Cactus (v3.0.1) (82). The resulting hierarchical alignment (HAL) file was converted to a multiple alignment file (MAF) with *M. lewisii* LF10 as the reference using cactus-hal2maf (83).

### Molecular evolutionary analysis of TERT

From the genome alignment we extracted the TERT multispecies alignment using the program maf_sort from the multiz package (84). Using the TERT multispecies alignment, the PAML codon substitution models (31) M0, M1a, M2a, M2arel, M3, M7, M8a, M8, and clade model C (32, 85) was fit using the program EasyCodeML (86). Evidence of relaxation of selection in the TERT alignment data was estimated using the method RELAX (33) implemented in the online Datamonkey server (87).

### Constructing plasmids for yeast three-hybrid analysis

We obtained the plasmids pIIIA/MS2-2 (Addgene plasmid #220632) and pGADT7 (Takara catalog #630442) to generate the vectors for yeast three-hybrid analysis. The full length sequences of telomerase RNA (TR) were PCR amplified from *M. lewisii*, *M. verbenaceus* and *M. cardinalis* and cloned into the *Smal* site of the pIIIA/MS2-2. In the pGADT7 plasmid the Gal4 transcription activation domain was fused to the TRBD region (position 229-580 of *M. lewisii* TERT) and cloned into the *NdeI* site. Constructs for the domain swap experiment were generated by switching the N-terminal region of *the* TRBD (position 229-386) with the C-terminal region (position 387-580) between *M. lewisii* and *M. verbenaceus*. For the single amino acid substitution constructs, the *M. lewisii*-specific residues lysine (K) at position 277 and tyrosine (Y) at position 309 were substituted with the corresponding *M. verbenaceus* residues glutamine (Q) and cysteine (C), respectively. Three constructs were generated containing the individual substitutions K277Q and Y309C, as well as the double substitution K277Q+Y309C. All amino acid substitution inserts were cloned into the *NdeI* site of the pGADT7 plasmid.All the primers used in this study are listed in Supplemental Table 7. Plasmids were ordered and generated by GenScript and vector sequences were validated by sequencing the whole plasmid with high coverage using nanopore sequencing serviced by Plasmidsaurus.

### Conducting yeast three-hybrid analysis

The Gal4 and MS2 constructs were transformed into yeast strain YBZ-1, which has the genotype *MATa, ura3-52, leu2-3, 112, his3-200, trp1-1, ade2, LYS2 :: (LexAop)-HIS3, ura3 :: (lexA-op)-lacZ, LexA-MS2 coat (N55K)*. Transformation was conducted following standard yeast two-hybrid protocol (88). After transformation selection was performed on media lacking leucine and adenine (–LA). The TERT and TR interactions were assessed by monitoring growth of YBZ-1 cells on selection plates lacking leucine, adenine, and histidine (–LAH). In addition, the strength of the TERT and TR interaction was assessed using 1mM, 3mM and 5mM of 3-amino triazole (3AT) concentrations. Each construct combination was co-transformed three times, followed by a minimum of three separate drop assays.

### Alphafold protein folding analysis

The *M. lewisii* TERT TRBD model was generated using AlphaFold3 (89). The model was analysed and visualized using UCSF ChimeraX (90).

### Combining TRAP with NGS sequencing (TRAP_NGS_)

Apical meristem were collected from three different plants of *M. lewisii, M.cardinalis*, and *M.verbenaceus* and immediately frozen in liquid nitrogen. Protein extraction and telomerase repeat amplification protocol (TRAP) assay were performed as previously described for *Mimulus* (22). The primers used for TRAP were previously tested and shown to work in the Erythranthe group *Mimulus* (22), and these were TS21 as the substrate primer for the telomerase and HisPRlong as the reverse primer for the PCR amplification (see Supplemental Table7 for information on primers).

50μL of TRAP-PCR product was used for preparing Illumina sequencing libraries. Libraries were prepared using the NEBNext® Ultra™ II DNA Library Prep Kit for Illumina® (NEB #E7645S) according to the manufacturer’s instructions. We conducted a PCR enrichment of adapter-ligated libraries using 7 PCR cycles to reach the required minimal concentration for sequencing. After construction of the library we took 5μl in volume from each library we made and pooled it together for Illumina sequencing. For the experiment testing the different TRAP PCR cycles libraries from the 4 different PCR cycles were made for sequencing libraries and pooled together for sequencing. For the experiment testing telomerase activity between species, the TRAP products from the 3 species with the 3 biological replicates were made into sequencing libraries and pooled together for sequencing. Pooled libraries were sequenced using the Illumina MiSeq Micro platform with PE-150 conformation.

### TRAP_NGS_ data analysis

Raw FASTQ data from TRAP_NGS_ were preprocessed before any downstream analysis using bbduk from the bbtools package (91). We used the sequences of TS21 and HisPR to bioinformatically cutoff the primers from each TRAP sequencing reads, and selected to analyze reads that were less than 96 bp to avoid analyzing reads that were not generated by the TRAP assay. Postprocessed sequencing reads were decomposed into the underlying telomere k-mer using TRviz (v1.2.0) (92). We used the sequence TTTCGG or TTTCGGG as the k-mer for the decomposition and counted the number of telomere repeats for each sequencing read. We assumed each sequencing read was an end product of a telomerase and classified if a sequencing read was a product of using the TR1 or TR2 molecule. This was accomplished by creating a sequence of perfect TTTCGG or TTTCGGG repeat and the one with the least amount of nucleotide differences to the observed sequencing read was chosen as the telomere repeat k-mer. We used the SequenceMatcher function from the Python (v3.12.0) difflib package to count the number of mutations. The script that implements our approach and classifies each sequencing read as a TTTCGG or TTTCGGG repeat is available at https://github.com/jychoilab/MimulusTERTevo.

### Bacterial DNA spike-in TRAP_NGS_ analysis

To validate our TRAP_NGS_ results and enable quantitative comparisons among *M. cardinalis*, *M. lewisii*, and *M. verbenaceus*, we repeated the TRAP_NGS_ experiment using an internal *Escherichia coli* genomic DNA spike-in. Sample collection, protein extraction, and the TRAP assay were performed as described above, except genomic DNA from *Escherichia coli* were included as spike-in before library preparation. Genomic DNA was prepared from NEB® 5-alpha Competent *E. coli* cells (High Efficiency; New England Biolabs, C2987H) and DNA was extracted using the Wizard® HMW DNA Extraction Kit (Promega).

To generate DNA fragments comparable in size to the TRAP amplification products, the *E. coli* genomic DNA was sheared using a Covaris M220 Focused-ultrasonicator. Sonication was performed in microTUBE-50 AFA Fiber Screw-Cap tubes (Covaris, PN 520166) using the M220 Holder XTU (PN 500414) with the corresponding XTU Insert for microTUBE 50 μL (PN 500488). Samples were processed at 20°C using the following settings: peak incident power of 75 W, duty factor of 10%, cycles per burst of 200, and treatment time of 1,600 sec. The sonication procedure was repeated three consecutive times on the same DNA sample to obtain a fragment size distribution centered at approximately 120 bp, closely matching the size of the TRAP amplification products. The fragment size distribution of the sheared *E. coli* genomic DNA was verified using an Agilent 4200 TapeStation prior to library preparation. A total of 5 ng of sheared *E. coli* genomic DNA was added to 50μL of TRAP-PCR product for each sample before library preparation. The library preparation and sequencing was performed as described in the previous section except that each library was individually quantified, normalized to an equal molar concentration, and pooled to ensure equal representation of each sample in the final sequencing library.

Raw Illumina reads from TRAP_NGS_ were first processed to remove adapter sequences and low-quality bases using bbduk from the bbtools package (91). The processed reads were aligned to the *E. coli* K-12 MG1655 reference genome (NCBI accession NC_000913.3) using BWA-MEM2 v2.2.1 (93) with default parameters. Reads mapped to the bacterial genome were used to estimate spike-in genome coverage using SAMtools v1.21 (94). The unmapped reads, representing TRAP assay products, were extracted and analysed as described in the previous section. In the end, the telomerase productivity represents the number of TRAP products per 1x *E. coli* genome read depth.

### Generating seedling and apical meristem transcriptomes for four *Mimulus* species

Tissues from two developmental stages were grown and harvested from *M. lewisii*, *M. cardinalis*, *M. verbenaceus*, and *M. parishii* for the transcriptome analysis. First we harvested above ground shoots from 14-day-old seedlings, and second we harvested apical meristem (AM) from 6 week old plants. For the seedling a biological replicate constituted roughly a dozen plants. For AM we collected the upper most tip of a plant and combined six to seven plant tissues as a biological replicate. We aimed to collect four biological replicates for each species–tissue combination, except for *M. lewisii* AM for which three biological replicates were obtained. Immediately after collection all samples were flash frozen in liquid nitrogen and stored at −80°C until RNA extraction. RNA was extracted using TRIzol (Invitrogen) according to the manufacturer’s protocol. The RNA was further treated with DNaseI (New England Biolabs) followed by column purification using Monarch Total RNA Miniprep Kit (New England Biolabs). Sequencing library was made using NEBNext Stranded mRNA Library Prep kit and sequenced using NextSeq2000 P3 single end 100bp read mode.

### Transcriptome data analysis

Raw RNA-seq reads were quality assessed using FastQC v0.12.1 (http://www.bioinformatics.babraham.ac.uk/projects/fastqc). Adapter sequences and low-quality bases were removed using fastp v0.24.0 (95). Filtered reads were then aligned to the corresponding *Mimulus* reference genomes using HISAT2 v2.2.1 (96), and resulting BAM files were sorted and indexed using SAMtools v1.21 (94). Gene-level read counts were generated using featureCounts v2.0.8 from the Subread package (97).

Analysis of gene expression differences between *Mimulus* species were conducted by first establishing the orthology between genes. We used Orthofinder v3.1.5 (98) to determine the orthogroups of the proteome sequences from the four *Mimulus* species (*M. cardinalis*, *M. lewisii*, *M. parishii*, and *M. verbenaceus*). Proteome sequences from evolutionarily distant plant species (*Aquilegia coerulea* (basal eudicot), *A. thaliana* (rosid), *Nymphaea colorata* (basal angiosperm), *Oryza sativa* (monocot), *Zea* mays (monocot)) were obtained from Phytozome (99) and were used as an outgroup for further parsing the *Mimulus* orthogroups into Phylogenetic Hierarchical Orthogroups (100), which uses gene based phylogenetic trees to further determine the orthology of duplicated genes. We focused on orthogroups that had a 1 to 1 orthology among all four *Mimulus* species. A *N* x *M* gene expression matrix was generated, where *N* represent the number of orthogroups and *M* represent the number of RNA-seq libraries for all four *Mimulus* species, which was loaded into edgeR v4.4.0 (101) for downstream transcriptome analysis. Lowly expressed orthogroups were filtered using filterByExpr() and retained counts were normalized using the trimmed mean of M-values (TMM) method.

Principal component analysis (PCA) was performed using the normalized log CPM expression matrix to evaluate transcriptomic relationships among species and tissue types. Differential expression analysis was performed using a generalized linear model with a factorial design: Expression ∼ Species × Tissue. Dispersions were estimated from biological replicates, and differential expression was assessed using the quasi-likelihood F-test (QLF) framework. Statistical significance was determined using Benjamini–Hochberg false discovery rate (FDR) correction, with orthogroups considered differentially expressed when FDR < 0.05 and absolute log_2_ fold change > 1.

### Telomere maintenance gene expression analysis

We focused our gene expression analysis to a curated set of 16 genes with previous experimental evidence of involvement in telomere maintenance and chromosome end protection in plants. Only telomere maintenance genes where we could establish 1 to 1 orthology among all four *Mimulus* species were examined for downstream analysis. These genes were CTC1 (102), KU70 (103), KU80 (104), MRE11 (105), NBS1 (106), NOP2A (107), POT1B (108), RAD50 (109), RPA70A (110), RPL5A (107), STN1 (111), TAC1 (112), TERT (45), TRB1 (113), TRB4 (113), and WHY1 (114). We used orthology with *A. thaliana* to determine which *Mimulus* genes corresponded to the telomere maintenance genes. During the orthology analysis we found POT1B had undergone a *Mimulus* specific duplication and we labeled the duplicates as POT1B-1 and POT1B-2. Normalized log CPM expression values corresponding to these genes were extracted from the ortholog-based expression matrix for downstream analyses. For visualization, biological replicates were merged by calculating the mean normalized log CPM expression value for each species–tissue combination. Gene expression values were subsequently standardized using row-wise z-score transformation to facilitate comparison of relative expression patterns among tissues and species.

### Telomere long read analysis of F1 hybrids between *M. lewisii* and *M. verbenaceus*

F1 hybrids were generated by crossing *M. lewisii* and *M.verbenaceus* and nanopore sequenced using the same protocols as the comparative genomic analysis. For the parental *M. lewisii* and *M.verbenaceus* raw nanopore signal intensity data from our previous study (22) was used to re-basecall using dorado v0.9.6 duplex mode with super-accurate model. For the long reads from the hybrid to identify *M. lewsii* and *M. verbenaceus* specific telomere long reads the whole genome sequencing nanopore long reads were aligned to the *M. lewisii* and *M.verbenaceus* reference genomes using minimap2 version 2.1.1-r341(115). The reference genomes are chromosomal level genome assemblies and reads that aligned to the first and last 10 kbp of each chromosome were extracted as these correspond to the putative telomere long reads.

The putative telomere long reads were filtered to select for true telomere reads using Topsicle (29). Topsicle was used to evaluate each putative telomere long read at the first and last 2000 bp to calculate the proportion of telomere repeats (*i.e.* the TRC parameter in Topsicle). Reads with TRC statistics greater than 0.85 were deemed a true telomere long read.

Analysis was performed on the FASTQ file containing uniquely aligned reads to the M. lewisii genome (F1_Lew_) and the *M. verbenaceus* genome (F1_VER_). Scripts used for visualizing the repeat distribution are available at https://github.com/jychoilab/MimulusTERTevo.

## Data availability

Nanopore sequencing data is available at the Sequence Read Archive (SRA) bioproject ID PRJNA1049363 under SRR38942558-SRR38942561 for *M. lewisii* LH2, SRR38942562 and SRR38942563 for *M. lewisii* JAC2, SRR38942564-SRR38942565 and SRR38942576-SRR38942577 for *M. lewisii* PPK, SRR38942566-SRR38942569 for *M. platycalyx*, SRR38942570 and SRR38942571 for *M. rupestris*, SRR38942572 and SRR38942573 for *M. flammea*, SRR38942574 and SRR38942575 for *M. cinnabarina*. Illumina sequencing data generated for TRAP sequencing is available under SRR39881933, SRR39881956, SRR39881957, and SRR39881958 for experiments testing PCR cycle effects, SRR39881934-SRR39881939, SRR39881948, SRR39881959, and SRR39881960 for testing telomerase activity, and SRR39881940-SRR39881955 for testing telomerase activity with spike in bacterial DNA. The comparative transcriptomic data is available under SRR39688617-SRR39688647. The nanopore sequencing data for conducting the F1 hybrid telomere sequence analysis was deposited under XXX. Raw data generated during the analysis including the genome assemblies, genome alignments, TERT multispecies alignment, and gene transcript counts are available at zenodo https://doi.org/10.5281/zenodo.20341648. Bioinformatic scripts and commands used for the study are available at https://github.com/jychoilab/MimulusTERTevo.

## Acknowledgement

We thank Dr. Stanley Fields for sharing the YBZ-1 strain. We also thank Drs. Lila Fishman, John Kelly, Miguel Flores Vergara, and John Willis for sharing the *Mimulus* species and ecotypes. We thank Dr. Sarah Zanders for advice on conducting the yeast genetic experiments. This study was supported by the National Institute of General Medical Sciences (NIGMS) of the National Institutes of Health award number R35GM154595. Illumina sequencing was made possible by the services from KU Genome Sequencing Core supported by NIGMS award number P30GM145499.

