## Supplementary figures and images for "Rapid evolution and functional divergence of the monkeyflower *Mimulus lewisii* telomerase"

### Fig S1

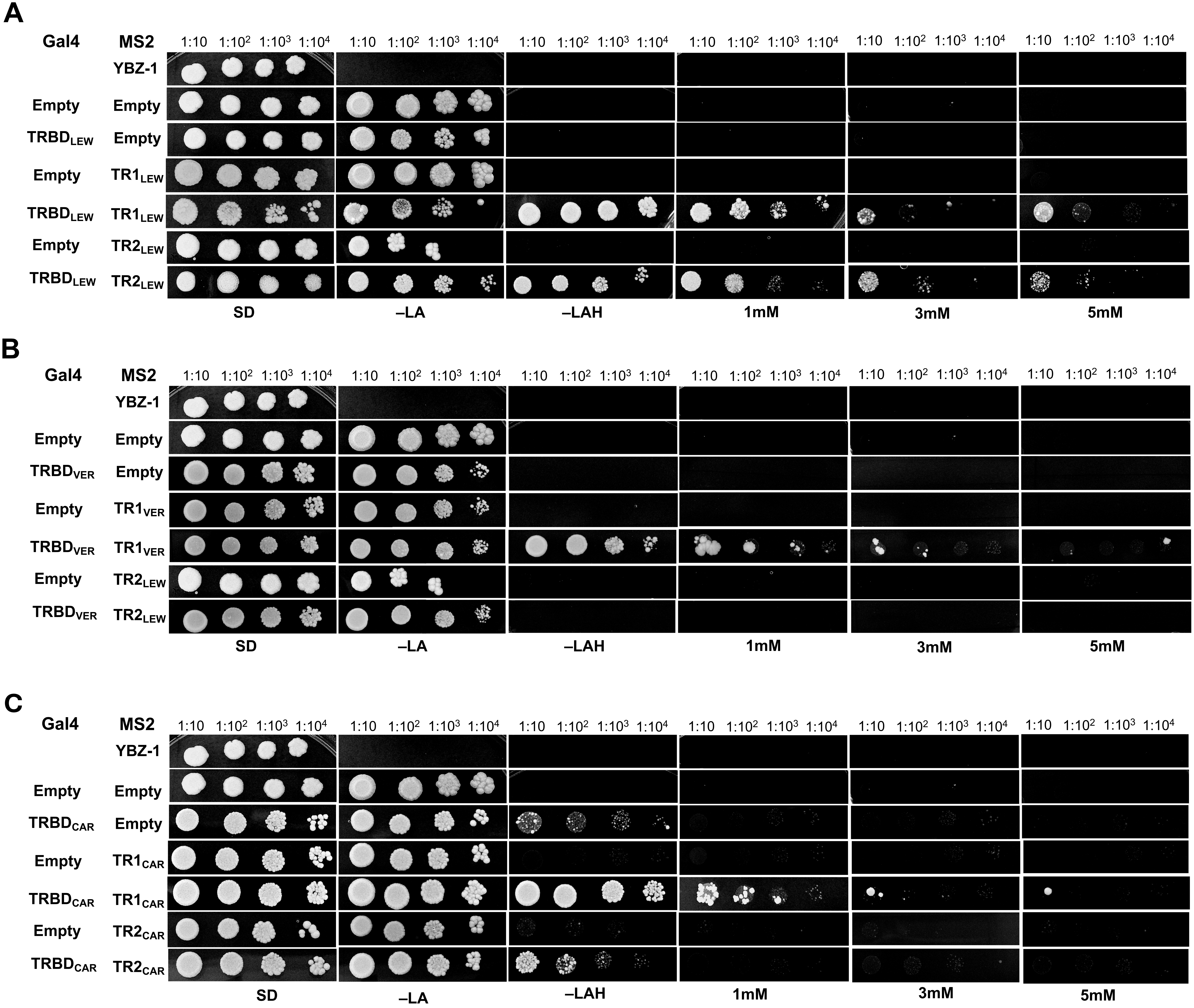

### Fig S2

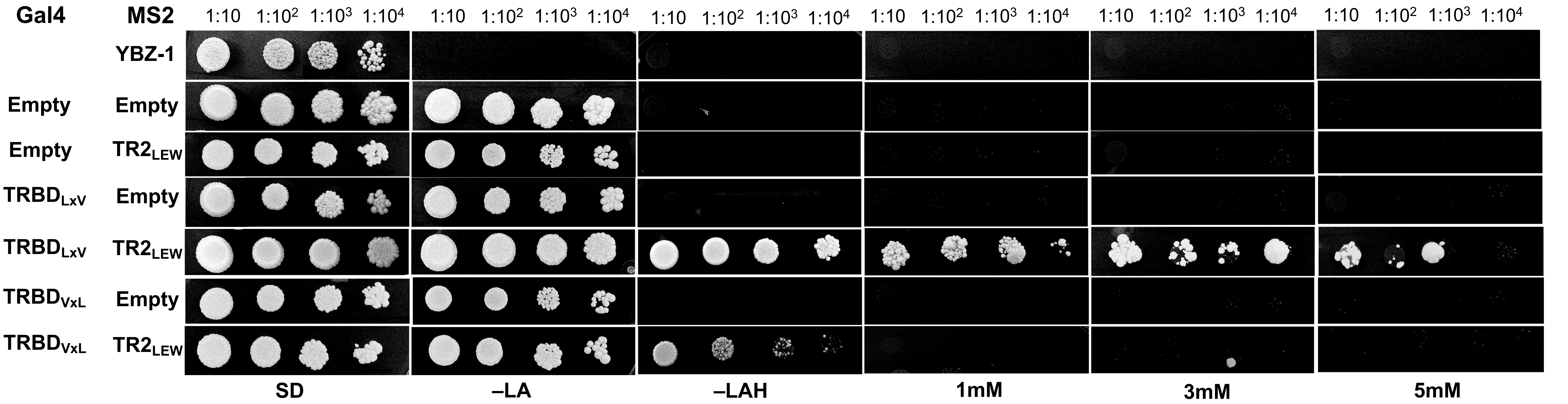

### Fig S3

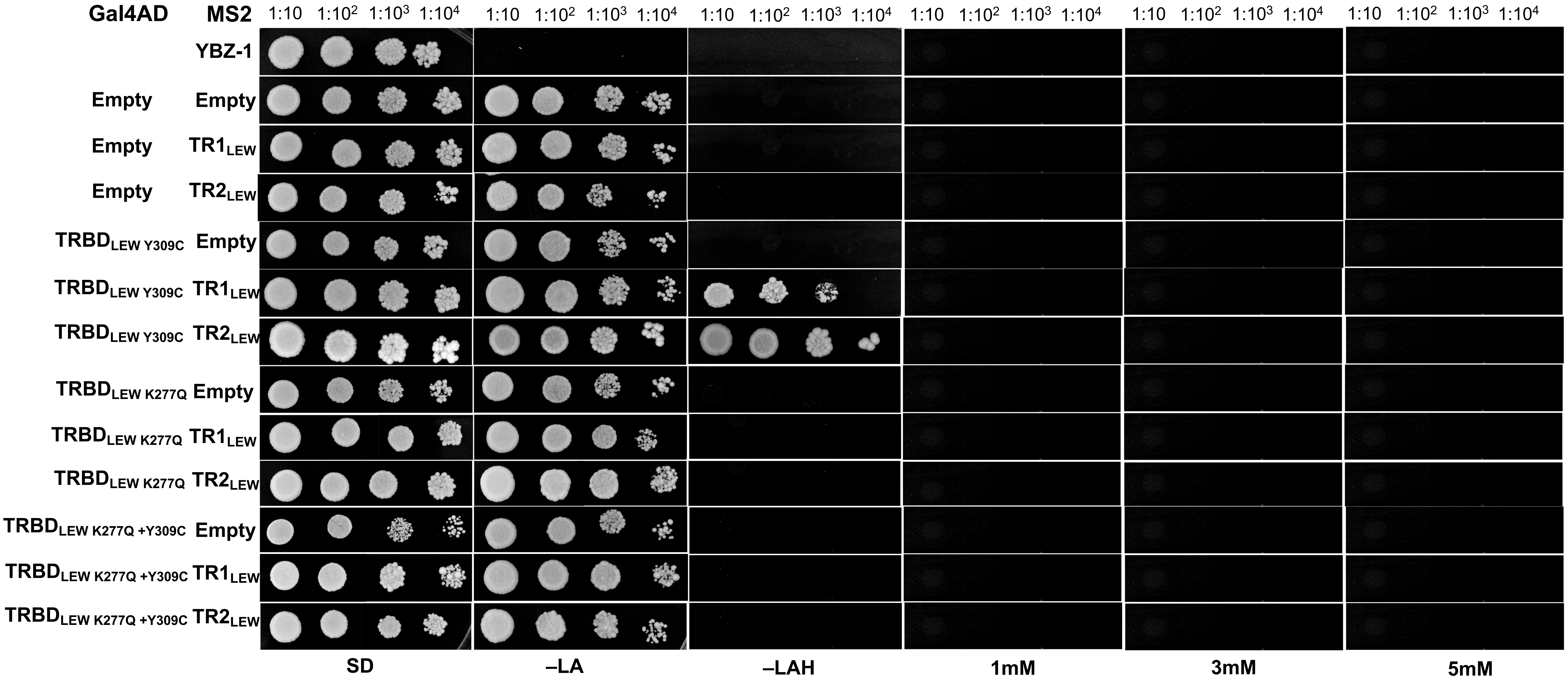

### Fig S4

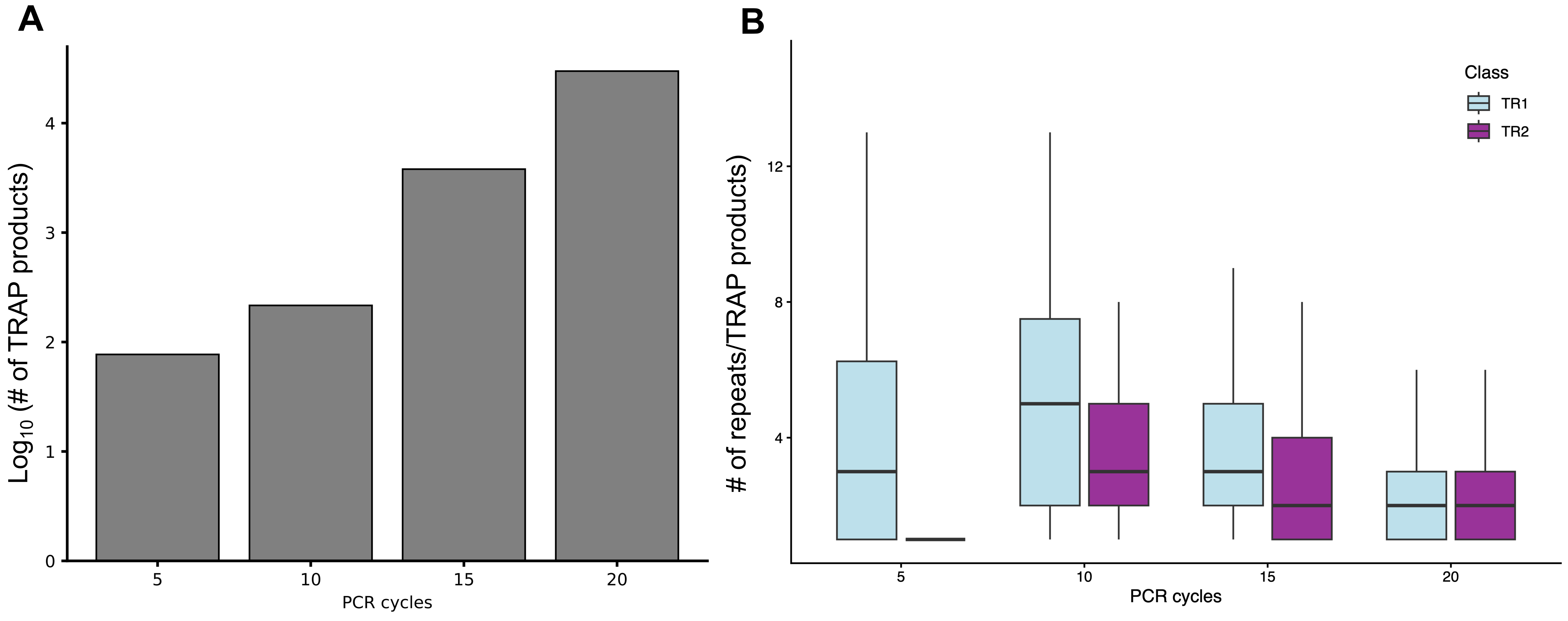

### Fig S5

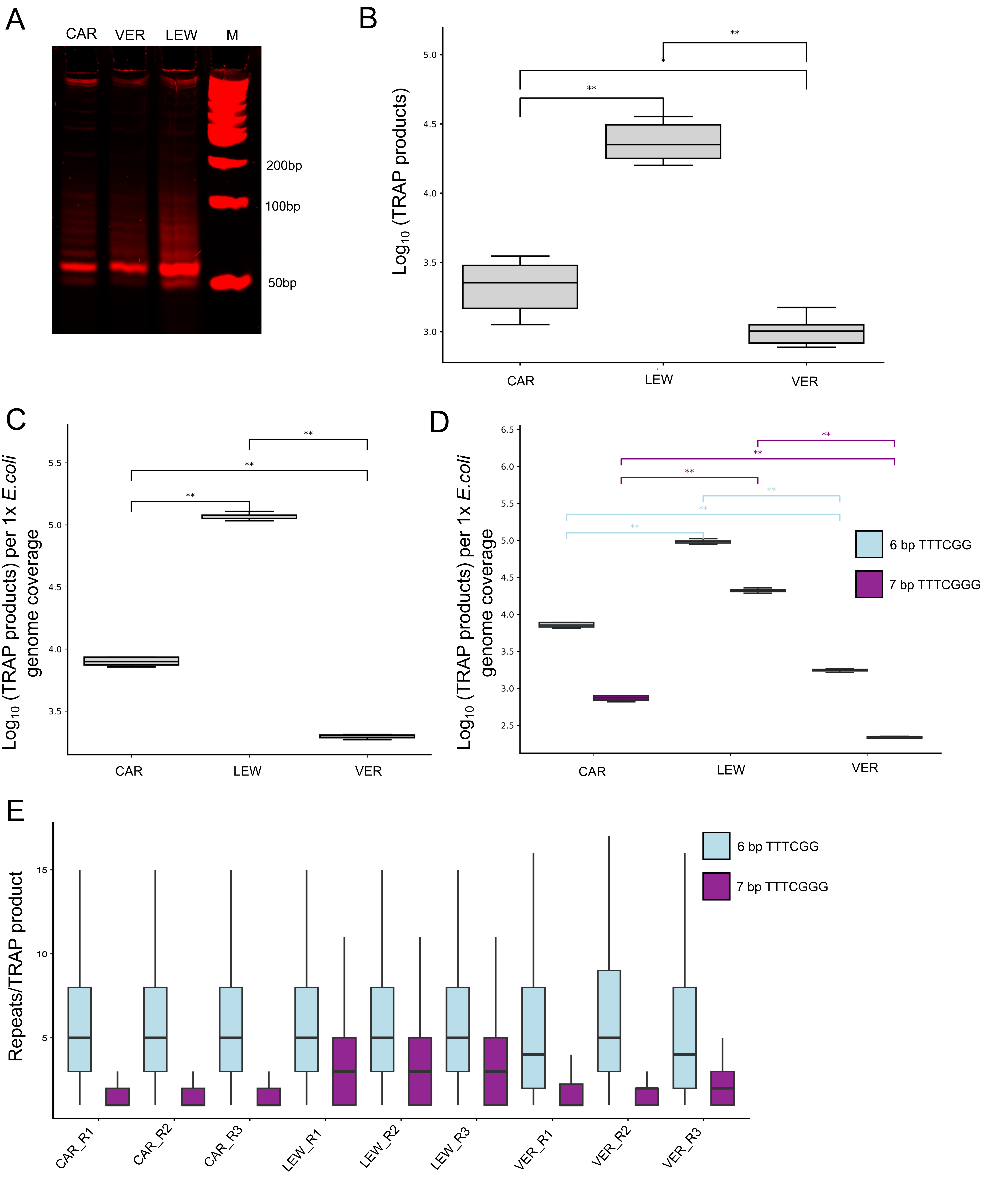

### Fig S6

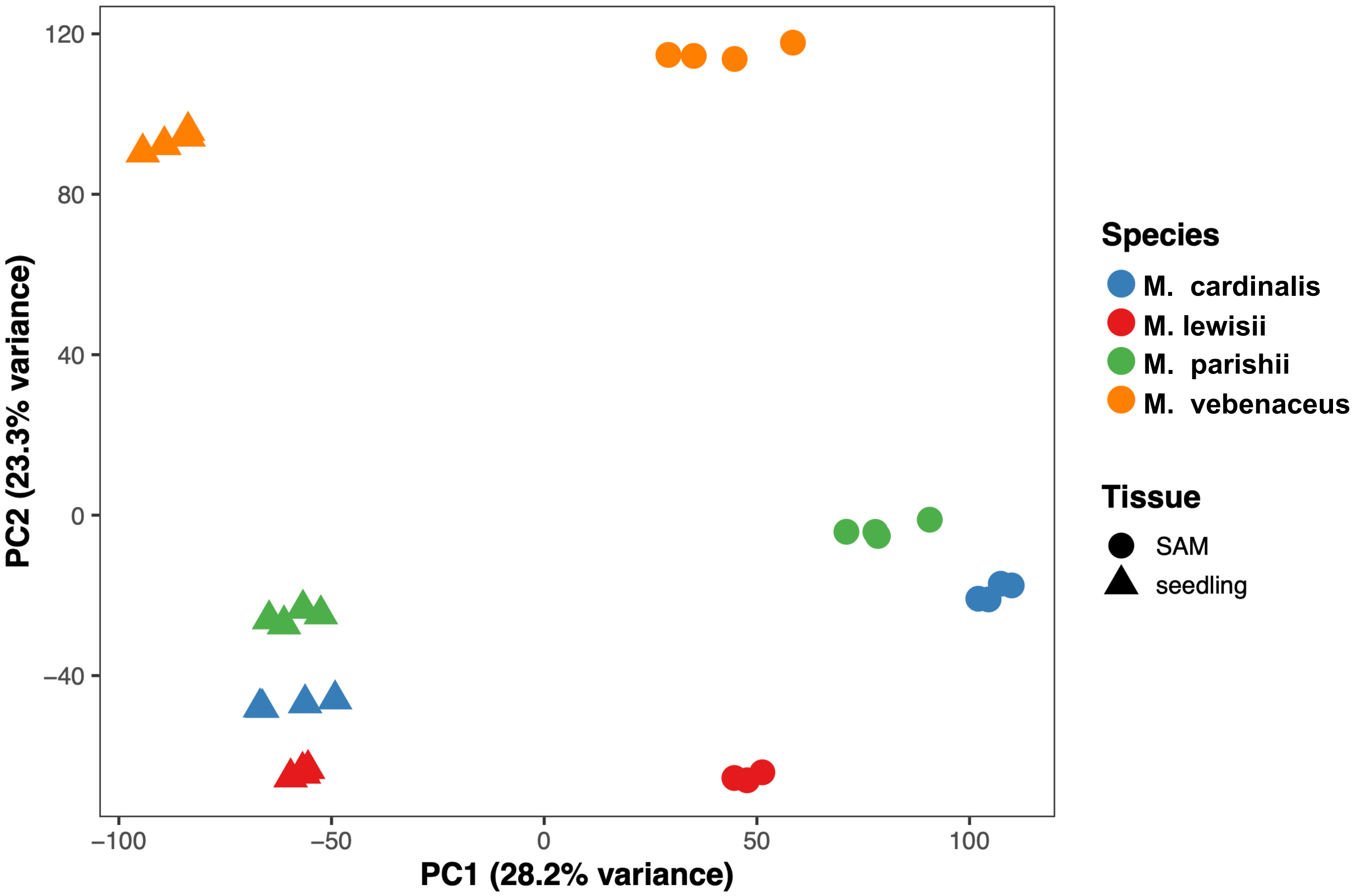

### Fig S7

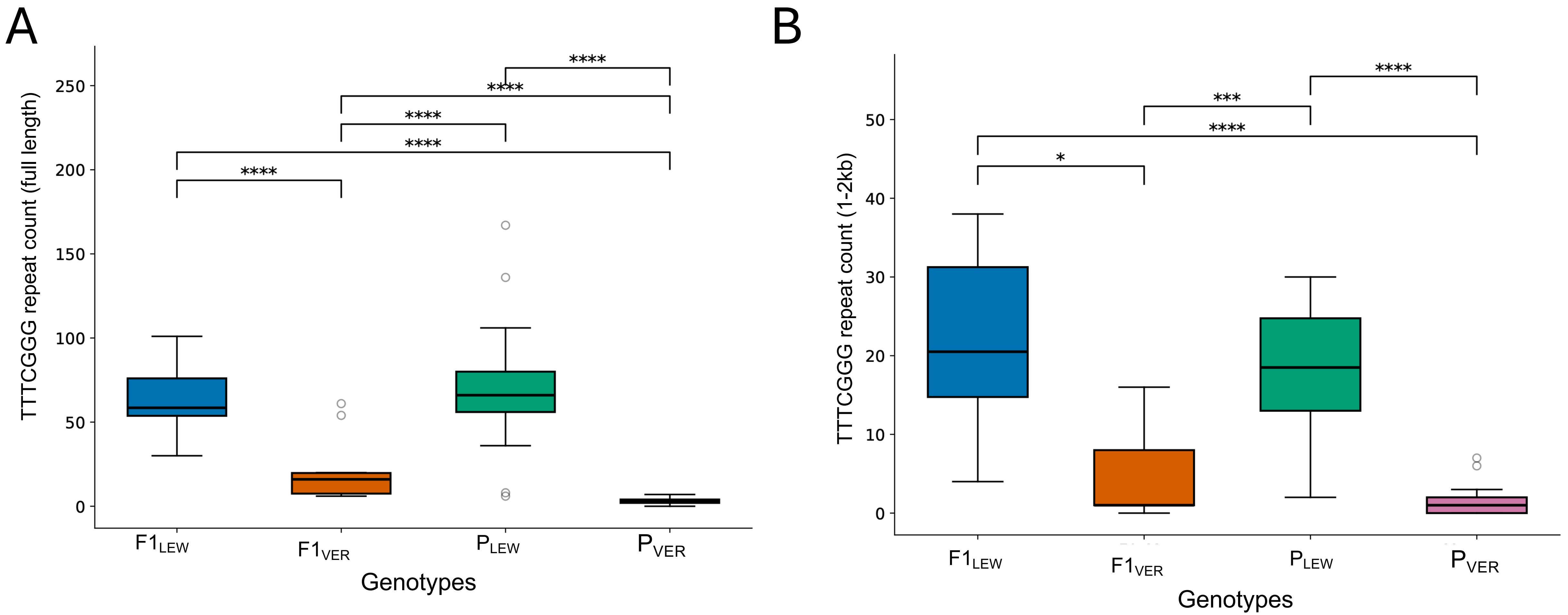
